# DeCiFering the Elusive Cancer Cell Fraction in Tumor Heterogeneity and Evolution

**DOI:** 10.1101/2021.02.27.429196

**Authors:** Gryte Satas, Simone Zaccaria, Mohammed El-Kebir, Benjamin J. Raphael

## Abstract

Most tumors are heterogeneous mixtures of normal cells and cancer cells, with individual cancer cells distinguished by somatic mutations that accumulated during the evolution of the tumor. The fundamental quantity used to measure tumor heterogeneity from somatic single-nucleotide variants (SNVs) is the Cancer Cell Fraction (CCF), or proportion of cancer cells that contain the SNV. However, in tumors containing copy-number aberrations (CNAs) – e.g. most solid tumors – the estimation of CCFs from DNA sequencing data is challenging because a CNA may alter the *mutation multiplicity*, or number of copies of an SNV. Existing methods to estimate CCFs rely on the restrictive Constant Mutation Multiplicity (CMM) assumption that the mutation multiplicity is constant across all tumor cells containing the mutation. However, the CMM assumption is commonly violated in tumors containing CNAs, and thus CCFs computed under the CMM assumption may yield unrealistic conclusions about tumor heterogeneity and evolution. The CCF also has a second limitation for phylogenetic analysis: the CCF measures the presence of a mutation at the present time, but SNVs may be lost during the evolution of a tumor due to deletions of chromosomal segments. Thus, SNVs that co-occur on the same phylogenetic branch may have different CCFs.

In this work, we address these limitations of the CCF in two ways. First, we show how to compute the CCF of an SNV under a less restrictive and more realistic assumption called the Single Split Copy Number (SSCN) assumption. Second, we introduce a novel statistic, the *descendant cell fraction* (DCF), that quantifies both the prevalence of an SNV *and* the past evolutionary history of SNVs under an evolutionary model that allows for mutation losses. That is, SNVs that co-occur on the same phylogenetic branch will have the same DCF. We implement these ideas in an algorithm named DeCiFer. DeCiFer computes the DCFs of SNVs from read counts and copy-number proportions and also infers clusters of mutations that are suitable for phylogenetic analysis. We show that DeCiFer clusters SNVs more accurately than existing methods on simulated data containing mutation losses. We apply DeCiFer to sequencing data from 49 metastatic prostate cancer samples and show that DeCiFer produces more parsimonious and reasonable reconstructions of tumor evolution compared to previous approaches. Thus, DeCiFer enables more accurate quantification of intra-tumor heterogeneity and improves downstream inference of tumor evolution.

**Code availability:** Software is available at https://github.com/raphael-group/decifer

## 1 Introduction

Cancer arises from an evolutionary process during which somatic mutations accumulate in the genome of different cells, yielding a heterogeneous tumor composed of different subpopulations of cells, or *clones*, that have distinct complements of mutations^1^. Quantifying the heterogeneity within a tumor is essential for understanding carcinogenesis and devising personalized treatment strategies^2–4^. While recent single-cell DNA sequencing technologies enable high-resolution measurements of tumor heterogeneity^5–11^, the vast majority of cancer studies in research and clinical settings^12–14^ rely on DNA sequencing of bulk tumor samples, where an individual sample comprises a mixture of thousands of different tumor cells. To quantify tumor heterogeneity using bulk sequencing data, most cancer sequencing studies analyze somatic single-nucleotide variants (SNVs) as these mutations are ubiquitous in cancer. The fundamental quantity used to quantify tumor heterogeneity^13,15,16^ from SNVs is the *Cancer Cell Fraction (CCF)* – also known as the *cellular prevalence* or the *mutation cellularity* – which is the proportion of cancer cells that contain the SNV. CCFs form the basis for many cancer analyses including: studying tumor heterogeneity^13,17-19^, reconstructing clonal evolution and metastatic progression13, identifying selection^20–22^, and analyzing changes in mutational processes over time^23–25^. In these and other studies, the underlying assumption is that groups of SNVs with the same CCF are likely to be present in the same cancer cells and thus occurred on the same branch of the phylogenetic tree that describes the evolution of the tumor (Figure 1a).

**Figure 1:**
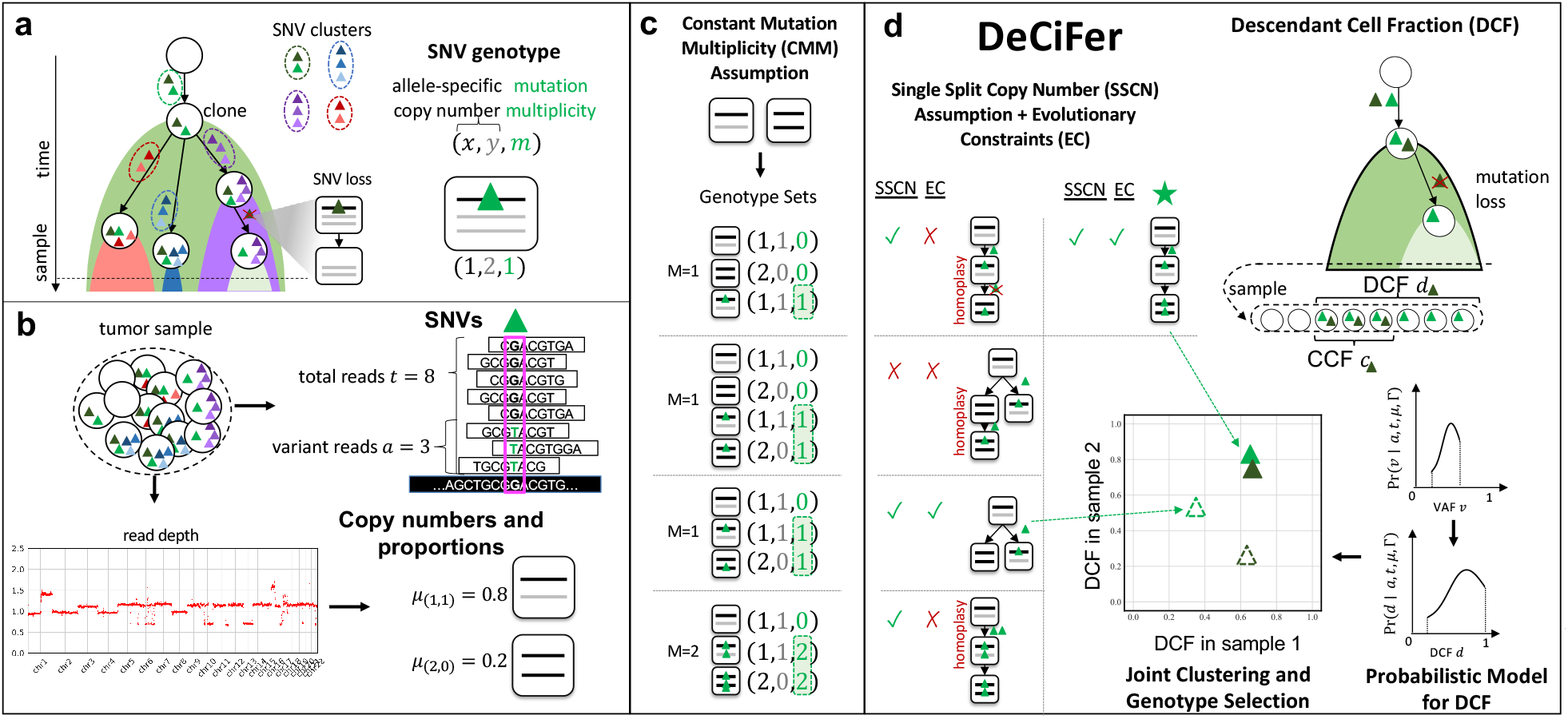
DeCiFer accurately groups SNVs into clusters with shared evolutionary history, accounting for the occurrence of copy number aberrations and mutation loss. **a**, During tumor evolution, SNVs in the same clone prior to a clonal expansion result in clusters of SNVs that are present in the same cells. Deriving these SNV clusters aids in identifying the clones in the tumor at the present time and in inferring the past evolutionary history of the tumor. CNAs may overlap SNVs and change their mutation multiplicities including losses of SNVs. The genotype of an SNV in a cell is composed of the copy number at the locus and the mutation multiplicity of the SNV. **b**, DNA sequencing of a bulk tumor sample yields two signals: (Right) the number of total and variant sequencing reads per SNV locus, which can be used to deduce the variant allele frequency (VAF) of the SNV, and (Bottom) the read depth across genomics regions which can be used to infer copy numbers and their proportions at SNV loci. **c**, The CMM assumption is used by nearly all existing methods to enumerate potential genotypes. The CMM assumption produces genotype sets where all cell genotypes that contain the mutation have the same number *M* of copies of the mutation (highlighted in green). **d**, DeCiFer uses a less restrictive assumption on genotypes, the SSCN assumption, as well as evolutionary constraints to enumerate potential genotype sets. DeCiFer excludes CMM genotype sets that are biologically unlikely (red crosses) but includes additional genotype sets (green star) that do not have constant mutation multiplicities. DeCiFer also computes the Descendant Cell Fraction (DCF) of each SNV, a statistic that summarizes both the prevalence of the SNV and its evolutionary history, and allows for mutation losses. DeCiFer simultaneously selects a genotype set for each SNV and clusters all SNVs using a probabilistic model of the DCF (or CCF).

Importantly, the CCF of an SNV is not directly measured in bulk DNA sequencing data of a tumor sample. Rather, the CCF must be inferred from the DNA sequencing reads that align to the locus containing the SNV. Specifically, the CCF is calculated from the *variant allele frequency* (VAF), or proportion of copies of the locus in the sample that contain the SNV. The VAF in turn is estimated as the proportion of variant reads at the SNV locus (Figure 1b). For a heterozygous SNV in a diploid genomic region, the CCF is twice the VAF. However, copy-number aberrations (CNAs) or loss-of-heterozygosity (LOH) events that overlap at an SNV locus can alter the number of copies of the SNV in a cell, and these events substantially complicate the estimation of the CCF. The reason for this added complexity is because the calculation of the CCF from the VAF depends on the *mutation multiplicity*, or the number of copies of the SNV in a tumor sample or clone. However, in bulk sequencing data only estimates of the total number of copies of a locus can be obtained^26–32^. Knowledge of the copy numbers at a locus – even allele-specific copy numbers and subclonal copy-number proportions – is insufficient to determine mutation multiplicities. Indeed, there are often multiple possible values for the unobserved mutation multiplicities that provide equally plausible explanations for the observed read counts and copy numbers at an SNV locus. In statistical terms, the CCF is not *identifiable* from DNA sequencing data (Figure S1a). Since CNAs and LOH events that amplify or delete large genomic segments, chromosomal arms, and even the whole genome^33–36^ are frequent in cancer – particularly in solid tumors where up to ∼90% of tumors^37^ may contain CNAs – it is imperative to have robust methods to calculate CCFs from bulk sequencing data.

Multiple computational methods have been developed in recent years to estimate CCFs in bulk sequencing data. These methods can be categorized into two different strategies. The first strategy is to severely restrict the possible mutation multiplicities ^13,16,21,35,38-45^ of SNVs; specifically, many methods assume that all cells harboring an SNV have the *same* mutation multiplicity. We refer to this assumption as the *Constant Mutation Multiplicity (CMM)* assumption (Figures 1c and S1b). Many methods rely on the CMM assumption, as well as additional heuristics, to select a single value of the CCF for each SNV. These methods either calculate the CCF for each SNV separately ^13,16,35,40,41,13^ or simultaneously infer CCFs and cluster SNVs across one or multiple samples, as done by PyClone^38^, Ccube^46^, and others ^39,44,47^. While the CMM assumption reduces the ambiguity in the calculation of the CCF, the assumption alone is insufficient to fully resolve such ambiguity (Figures 1c and S1a). Moreover, the heuristics used to select the mutation multiplicity of an SNV – e.g., rounding the estimated average mutation multiplicity to the nearest integer^41^ – may introduce unexpected biases into the resulting analyses. More importantly, the CMM assumption is often violated in practice (Figure S1b). For example, an SNV occurring before an amplification may result in cancer cells with different mutation multiplicities: a group of cells without the amplification and with a single copy of the SNV, and another group of cells with the amplification and multiple copies of the SNV. Scenarios such as this are frequent in solid tumors that often have subclonal CNAs ^12,27,33,36^. Thus, the CMM assumption is both too restrictive to model many real tumors and also too weak to overcome the issue of non-identifiability.

The second strategy to estimate CCFs is a phylogenetic approach using an evolutionary model that includes both SNVs and CNAs. Methods that use this strategy include PhyloWGS^48^, SPRUCE^49^ and Canopy^50^. The evolutionary models employed in these methods do not make the CMM assumption and thus allow more realistic scenarios such as mutation losses. However, this flexibility comes at a cost of computational efficiency: none of the current methods scale to the large numbers of SNVs measured in current cancer sequencing studies, and these methods may take days or weeks to run even samples with a relatively small number of mutations (≈ 1000). To address scalability, these methods group mutations into clusters where all mutations in a cluster are assumed to occur on the same branch of the phylogenetic tree describing the evolution of the tumor. Specifically, PhyloWGS^48^ simultaneously clusters mutations and reconstructs a phylogeny, while Canopy^50^ and SPRUCE^49^ require mutation clusters in input. However, this pre-clustering approach is difficult because the CCFs of the mutations are not known in advance. If one clusters mutations using CCFs derived under the CMM assumption then the restriction on mutation multiplicities imposed by the CMM assumption reduces or eliminates the advantage of the phylogenetic approach. Because of these limitations, phylogenetic methods are not as widely used as methods that rely on the CMM assumption.

In addition to the difficulties in estimating the CCF, there is another important limitation of the CCF itself: in many cases, the CCF is *not* the correct quantity to use for phylogenetic analysis. Specifically, the CCF measures only the prevalence of a mutation in the tumor at the *present* time and does not necessarily provide complete information about the *past* history of the mutation. Two mutations that occurred during the same cell division may have very different CCF values if one of these mutations is later lost due to a deletion^45^ (Figure S1c). In this case, one mutation may have a high CCF – suggesting that the mutation occurred early in the evolution of the tumor – while the other mutation has a low CCF value, misleadingly suggesting that the mutation occurred late during evolution. In fact, mutation losses are common in cancers that contain many CNAs ^34,51,52^. Jamal-Hanjani et al.^13^ described this issue in the TRACERx sequencing study of non-small-cell lung cancer patients and proposed the “phyloCCF”, an *ad hoc* correction of the inferred CCF for SNVs in genomic regions affected by subclonal deletions. However, the phyloCCF still relies on the CMM assumption and thus models only some of the effects of CNAs on CCFs.

In this paper, we propose a new approach to analyze tumor heterogeneity and evolution in tumors that contain both SNVs and CNAs, addressing both limitations in current approaches to estimate CCF and limitations in the CCF quantity itself for phylogenetic analysis (Figures 1d and S1d-f). We first show how to compute the CCF under the *Single-Split Copy Number* (SSCN) assumption, an assumption that relies on standard evolutionary models for SNVs and CNAs and is less restrictive than the CMM assumption. We then introduce a novel statistic, the *descendant cell fraction (DCF)*, that generalizes the CCF to account for mutation losses. The DCF provides an elegant mapping between the quantities measured in bulk DNA sequencing data – CNAs and read counts of SNVs – and the evolutionary history of SNVs. Specifically, SNVs that co-occur on the same branch of the phylogenetic tree will have the same DCF. We utilize this mapping to derive a probabilistic model to estimate the CCF or DCF while accounting for uncertainties in DNA sequencing data. Finally, we address the issue of non-identifiability in the CCF and DCF by sharing information across multiple SNVs and samples. We implement our approach in an algorithm named DeCiFer. DeCiFer can be viewed as an intermediate between scalable approaches that compute CCFs using the restrictive CMM assumption without an evolutionary model and phylogenetic approaches that simultaneously model the evolution of all SNVs and CNAs but do not scale to large numbers of mutations. DeCiFer combines the advantages of both approaches (e.g., clustering of mutations and joint evolution of SNVs and CNAs) while avoiding some of their major limitations (e.g., CMM assumption and scalability). We show that that DeCiFer outperforms existing methods on simulated data. Finally, we use DeCiFer to analyze 49 metastatic prostate cancer samples^17^, and we show that DeCiFer infers DCFs that result in more realistic and more parsimonious evolutionary histories for these tumors compared to existing approaches.

## 2 Methods

### 2.1 The Cancer Cell Fraction: Current Approaches

The cancer cell fraction (CCF) *c* of an SNV is defined as the fraction of cancer cells in a sample that contain at least one copy of the SNV. The CCF is not directly observed from bulk data; rather, one observes the total number *t* of reads that align to the SNV locus and the corresponding number *a* of reads with the variant allele (Figure 1b). If the SNV locus is diploid (i.e., no CNAs), the standard approach ^13,16,19,21,35,38-45,47^ estimates the CCF *c* from the fraction 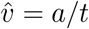 of variant reads as 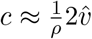, where ρ is the *tumor purity* – i.e., fraction of cancer cells in the sample – which also may be inferred from bulk data^26–32^. Note that 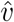 is the maximum likelihood estimate (MLE) estimate of the *variant allele frequency* (VAF) *v* – i.e., the proportion of copies of the locus in the sample that contain the SNV. More generally, if the SNV locus is aneuploid due to CNAs, nearly all existing methods ^13,16,21,35,38–45^ estimate the CCF by using the following generalization of the equation for diploid case:

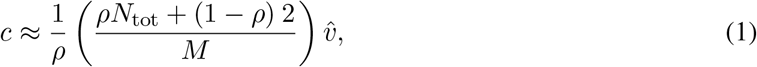

where *N*_tot_ is the average total copy number in cancer cells and *M* is the number of copies of the SNV in cancer cells that contain the SNV.

While Eq. (1) has become the standard in the field, this equation incorporates strong assumptions about tumor composition and evolution. To describe these assumptions, we define the *genotype* of an SNV locus in a cell as a triple (*x, y, m*) of non-negative integers, where the copy numbers (*x, y*) correspond to the number of maternal and paternal copies of the locus and the *mutation multiplicity m ≤ x* + *y* is the number of copies with the SNV. The CCF *c* is then the fraction of cancer cells that have genotypes (*x, y, m*) with *m* ≥ 1. Thus, we see that Eq. (1) assumes that *m* = *M* is fixed across all the cancer cells that contain the SNV (Figure 1c), which we state formally as follows.

#### Constant Mutation Multiplicity (CMM) Assumption

*At every SNV locus, there exists an integer M ≥* 1 *such that all genotypes at the locus have the form* (*x, y, m*) *where either m* = 0 *or m* = *M*.

### 2.2 The Cancer Cell Fraction: Single Split Copy Number Assumption

The CMM assumption severely limits the genotypes at a locus and is often violated in tumors with CNAs, as we will demonstrate in Results. Here, we define a less restrictive assumption on genotypes, the Single Split Copy Number (SSCN) Assumption, that facilitates the computation of CCFs from bulk sequencing data under commonly used evolutionary models. Formally, we define a *genotype set* Γ as the set of genotypes at an SNV locus. Each genotype (*x, y, m*) in Γ has a corresponding genotype proportion *g*_(*x,y,m*)_ that gives the prevalence of the genotype in the sample. Let 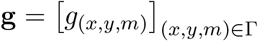 denote the genotype proportions for genotypes in Γ, and note that the genotype proportions satisfy *g*_(*x,y,m*)_ *≥* 0 and Σ _(*x,y,m*)∈Γ_ *g*_(*x,y,m*)_ = 1. Given tumor purity ρ and a pair (Γ, **g**) of a genotype set and genotype proportions the CCF *c* is *uniquely* determined by the following equation:

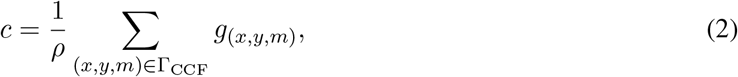

where Γ_CCF_ = {(*x, y, m*) ∈ Γ | *m ≥* 1} ⊆ Γ is the set of genotypes that contain the SNV.

Unfortunately, Eq. (2) is not directly applicable to bulk DNA sequencing data because neither the genotype set Γ nor the genotype proportions **g** are directly measured in bulk data. Rather from the aligned sequencing reads, one can estimate the VAF *v* of an SNV as well as the proportions *µ*_(*x,y*)_ of cells with copy number (*x, y*) at the locus. The copy-number proportions ***µ*** = [*µ*_(*x,y*)_] may be inferred using current tools for copy number deconvolution^26–32^. The key question is: what genotypes and genotype proportions are consistent with the estimated copy number proportions ***µ*** and VAF *v*? The relationship between these quantities is given by the following equations.

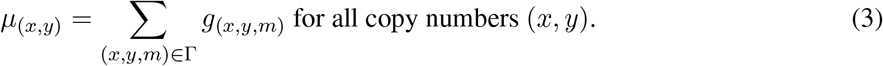

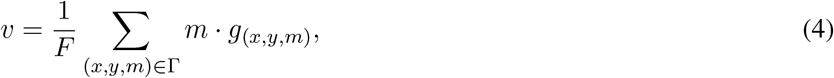

where *F* is the fractional copy number defined as Σ _(*x,y*)_(*x* + *y*) · *µ*_(*x,y*)_. Note that *F* is the average copy number over all cells, including both cancer and normal cells; in contrast *N*_tot_ in Eq. (1) is the average copy number in cancer cells only. Thus, we have that *F* = *ρN*_tot_ + 2(1 – ρ) for SNVs in autosomal chromosomes.

Given copy-number proportions ***µ*** and VAF *v*, there are often many pairs (Γ, **g**) that satisfy Eqs. (3) and (4); i.e., these equations are severely underdetermined. Thus, it is necessary to impose additional constraints on the pairs (Γ, **g**) that are considered. We make the following assumption.

#### Single Split Copy Number (SSCN) Assumption

*At every mutation locus, there is exactly one copy number* (*x*^*^, *y*^*^) *with two distinct genotypes* (*x*^*^, *y*^*^, 0) *and* (*x*^*^, *y*^*^, *m*^*^).

We call a genotype set Γ^*^ adhering to the SSCN Assumption an *SSCN genotype set*. SSCN genotype sets Γ^*^ have two desirable properties: (1) They arise from standard evolutionary models for SNVs and CNAs (see Section 2.5 for details); (2) If genotype proportions **g** satisfying equations Eqs. (3) and (4) exist, then they are unique (see Appendix Eq. (S5)). Note that these properties are not necessarily true for genotype sets derived from the CMM assumption. We say that a genotype set Γ is *consistent* with *v* and ***µ*** provided there exist corresponding genotype proportions **g** satisfying Eqs. (3) and (4). In Appendix B.1 we describe necessary and sufficient conditions for consistency.

As the genotype proportions **g** for an SSCN genotype set Γ^*^ are uniquely determined given VAF *v* and copy-number proportions ***µ***, the CCF *c* is uniquely determined as well (by Eq. (2)). Thus, we have the following relationship between CCF *c* and VAF *v* for an SSCN genotype set Γ^*^.

##### Theorem 1.

*Given tumor purity* ρ, *VAF v, copy-number proportions* ***µ***, *and an SSCN genotype set* Γ^*^ *consistent with v and* ***µ***, *the CCF c is uniquely determined by*

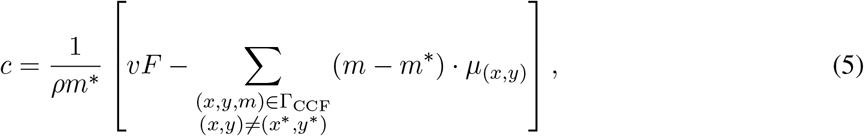

*where* Γ_CCF_ = {(*x, y, m*) ∈ Γ^*^ | *m* ≥ 1} ⊆ Γ^*^ *is the set of genotypes containing the mutation*.

Further details and the proof for Theorem 1 are in Appendix B.1. Note that similar to the CMM Assumption, the CCF is non-identifiable under the SSCN Assumption since ***µ*** and *v* are not sufficient to determine Γ^*^. We describe how to enumerate and select SSCN genotype sets Γ^*^ in Section 2.5.

### 2.3 Probabilistic model for the Cancer Cell Fraction

Recall that the VAF *v* is not directly measured from sequencing data; rather one observes only the total read count *t* and variant read count *a*. Thus, the affine transformation in Eq. (5) cannot be used directly to compute the CCF *c* from the VAF *v*. Many existing methods ^13,19,21,39–41,41,43,44^ calculate the CCF by assuming that the proportion 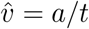 of variant reads is an accurate estimate of the VAF and do not evaluate uncertainty due to sequencing errors and coverage. Here we show how to derive a probability distribution Pr(*c*) for the CCF *c* from any probability distribution Pr(*v*) on the VAF *v*. Specifically, we compute the posterior probability Pr(*c* | *a, t*, ***µ***, Γ) of the CCF given the observed data and an SSCN genotype set as

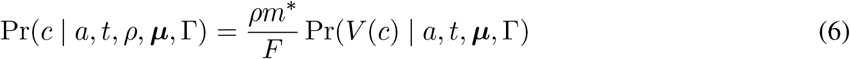

where 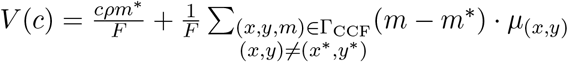 is the inverse of Eq. (5). We derive this formula using a change-of-variable technique in Appendix B.3.

To derive Pr(*V* (*c*) | *a, t*, ***µ***, Γ), we first apply Bayes’ Theorem, giving

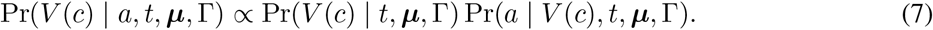

We then assume that the VAF is conditionally independent of the total read count *t* given ***µ*** and Γ and that variant read count *v* is conditionally independent of ***µ*** and Γ given the VAF *V* (*c*) and total read count *t*. This yields the following posterior probability for *V* (*c*).

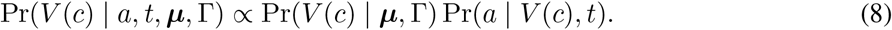

Pr(*a* | *V* (*c*), *t*) is the likelihood of the observed variant read counts for a given VAF value. Pr(*V* (*c*) | ***µ***, Γ) is a prior over VAFs. Pr(*V* (*c*) | ***µ***, Γ) is the prior probability of the VAF given copy-number proportion and a genotype set. In Appendix B.1, we describe that due to consistency constraints, not all VAFs are feasible given copy-number proportions ***µ*** and SSCN genotype set Γ. For example, 0.5 is an upper bound for VAFs *v* for heterozygous mutations in diploid regions. Thus, the prior for VAF *v* has support over the range [*v*^−^, *v*^+^] of feasible VAFs. One can use any reasonable distributions for the prior and likelihood. In practice, we use a uniform distribution over [*v*^−^, *v*^+^] for the prior, and a binomial or beta-binomial distribution for the likelihood. In Appendix B.3 we provide a more detailed derivation of the probabilistic model and in Appendix B.7 we describe how we estimate parameters for the beta-binomial distribution from DNA sequencing data.

### 2.4 Descendant Cell Fraction

We derive a new quantity, the *descendant cell fraction* (DCF), a generalization of the CCF that accounts for potential SNV losses. The DCF of a mutation is the proportion of cells in a sample that are descendants of the ancestral cell where the mutation was first introduced. As an example, consider two SNVs that occurred at the same time in the same cell. If one of these SNVs is subsequently lost due to a deletion, these SNVs would have distinct CCFs at the time of sampling. However, the DCF for both SNVs would be the same, as they have the same set of descendent cells in the sample. Note that the DCF equals the CCF for SNVs that are not affected by deletions.

To define the DCF formally, we introduce the notion of a *genotype tree T*_Γ_ = (Γ, *E*), which is a rooted tree whose vertex set is a genotype set Γ and whose directed edges *E* encode evolutionary relationships between pairs of genotypes. While a tumor phylogeny models the evolutionary history of all SNVs in the tumor, a genotype tree describes the evolutionary history of only a single SNV. As such, inference of genotype trees of individual SNVs is a less challenging task than comprehensive phylogeny inference. We summarize a genotype tree *T*_Γ_ and genotype proportions **g** by the DCF, which will enable us to assign a genotype tree to each SNV subject to a parsimony constraint regarding the number of distinct DCF values. Specifically, the DCF *d* of an SNV is defined as

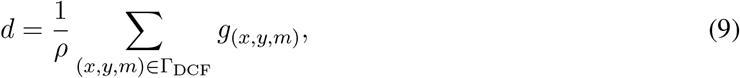

where Γ_DCF_ ⊆ Γ is the subset of genotypes that are *descendants* of the vertex corresponding to genotype (*x*^*^, *y*^*^, *m*^*^). Thus, while the CCF is the total prevalence of the subset Γ_CCF_ ⊆ Γ of genotypes with mutation multiplicity *m* 1 at the present time, the DCF is total prevalence of genotypes that are descendants of the genotype (*x*^*^, *y*^*^, *m*^*^) where the mutation is first introduced. We have the following theorem.

#### Theorem 2.

*Given tumor purity* ρ, *VAF v, copy-number proportions* ***µ***, *an SSCN genotype set* Γ^*^ *consistent with v and* ***µ***, *and a genotype tree T*_Γ_*, *the DCF d is uniquely determined by*

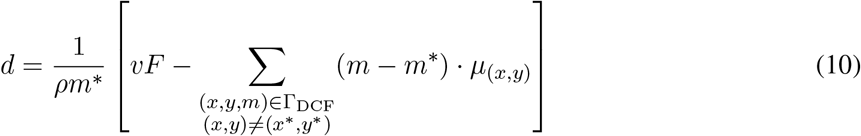

*where* Γ_DCF_ *is the set of genotypes in genotype tree T*_Γ_* *that are descendants of the state* (*x*^*^, *y*^*^, *m*^*^).

Observe that the only difference between Eqs. (10) and (5) is the use of Γ_DCF_ rather than Γ_CCF_. To obtain a probabilistic model for the DCF *d*, we follow a similar procedure described in Section 2.3 above for the CCF, replacing the inverse transformation *V*(*c*) with 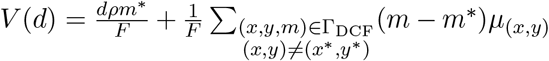 (see Appendix B.3).

### 2.5 DeCiFer: Simultaneous Clustering and Genotype Selection using the DCF

Theorem 2 defines DCF *d* given VAF *v*, copy-number proportions ***µ*** and a genotype tree *T*_Γ_* for SSCN genotype set Γ^*^. However, neither Γ^*^ nor *T*_Γ_* are directly observed by bulk data, and there frequently are multiple possible values that are consistent with the observed data (Figure S1a). Examining SNVs individually, there is no way to distinguish between these values. However, by evaluating SNVs jointly and assuming that there are a small number of possible values of the DCF, we obtain constraints that reduce ambiguity in the selection of Γ^*^ or *T*_Γ_* for individual SNVs. Specifically, we make the following assumption.

#### Assumption

*There exist DCF values d*_1_,..., *d*_*k*_ *such that for every SNV in a tumor sample at least one d*_*j*_ *is a valid DCF for the SNV (i*.*e*., *solution of equation* (10)).

Under this assumption, SNVs may be partitioned into *k* groups according to their DCF. However, since SNVs may have more than one possible DCF value, the problem of identifying these groups entails the simultaneous selection of a genotype tree for each SNV (which determines the DCF value of the SNV) and the clustering of SNVs into *k* groups according to their DCF values.

Here, we describe this simultaneous selection and clustering problem in the more general setting where we have observations from *p* bulk sequencing samples from the same patient. Thus, we are given variant read counts **a**_*i*_ = [*a*_*i,ℓ*_]_*ℓ*=1,…,*p*_, total read counts **t**_*i*_ = [*t*_*i,ℓ*_]_*ℓ*∈1,…,*p*_ and copy-number proportions *M*_*i*_ = [***µ***_*i,ℓ*_]_*ℓ*=1,…,*p*_ for each SNV *i* in each sample *ℓ*. Let *G*_*i*_ be the set of genotype trees *T*_Γ_* for SSCN genotype sets Γ^*^ that are consistent with ***µ***_***i***_. Let *s*_*i*_ ∈ {1,. .. |*G*_*i*_|} be a selection of a genotype tree *T*_Γ_* for SNV *i*. Let *z*_*i*_ ∈ {1,. .., *k*} be a cluster assignment for SNV *i*. We aim to find genotype selections **s**, cluster assignments **z**, and cluster DCFs *D* = [**d**_1_,. .., **d**_*k*_] where *d*_*j,ℓ*_ is the DCF of cluster *j* in sample *ℓ* that maximize the posterior probability of the parameters given the observed read counts. Eq. (S23) in Appendix B.3 gives this posterior probability of the DCF for an individual SNV in one sample. We assume that given cluster assignments and DCFs, variant read counts are conditionally independent across samples and across SNVs. Thus, to compute the objective, we take the product across samples and SNVs. This leads to the following problem.

##### Problem 1

(Probabilistic Mutation Clustering and Genotype Selection (PMCGS)). *Given a set 𝒢*_*i*_ *of pairs of genotype sets and trees, variant read counts* **a**_*i*_, *total read counts* **t**_*i*_ *and (iv) copy-number proportions* ***µ***_*i*_ *for each SNV i as well as an integer k >* 0 *find (i) DCFs* **D**^*^ = [**d**_1_,...,**d**_*k*_] *and for each SNV i, select (ii)* 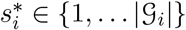 *and (iii)* 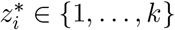 *such that*

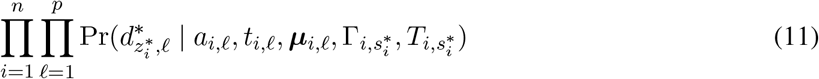

*is maximum*.

While the hardness of the PMCGS problem is open, the variant of the problem where every VAF *v* is observed instead of read counts *a, t* is NP-complete as it is equivalent to the well-studied HITTING SET problem (Appendix B.4).

We introduce DeCiFer, an algorithm to solve the PMCGS. DeCiFer uses a coordinate ascent approach to solve Eq. (11) by alternately optimizing (i) the cluster assignments **z** and genotype set assignments **s** for individual SNVs and (ii) the cluster DCFs **D**.

DeCiFer imposes stronger constraints on the allowed genotype sets than imposed by the SSCN assumption, since some SSCN genotype sets are not evolutionary plausible (for example, see the top left genotype set in Figure 1d). Specifically, we assume that the allowed genotype trees *T*_Γ_ for an individual SNV conform to the following evolutionary model. First, each mutation is introduced exactly once but may be subsequently lost or amplified due to CNAs. Second, each allele-specific copy number (*x, y*) of the SNV locus is attained exactly once. Thus, viewing SNVs as two-state characters and CNAs as multi-state characters, we have the Dollo model for SNVs and the infinite alleles assumption for CNAs. Finally, any change in mutation multiplicity must be caused by a corresponding change in copy-number. These constraints were formally described by El-Kebir et al.^49^ (Definition 12, Supplementary Material). Note that all genotype sets Γ that meet these constraints are also SSCN genotype sets. Under the evolutionary constraints, the mutation multiplicity *m*^*^ for the split copy number (*x*^*^, *y*^*^) is *m*^*^ = 1. DeCiFer enumerates *G*_*i*_ using the same tree enumeration procedure introduced in SPRUCE^49^.

Further details of DeCiFer model selection and implementation are in Appendix B.5. DeCiFer is available at https://github.com/raphael-group/decifer.

## 3 Results

### 3.1 Simulations

We assessed the performance of DeCiFer on simulated data with varying number of SNVs (100, 250, 500, or 1000), samples (1, 3, 5, or 7) and SNV clusters (2, 3, 5, or 8) as well as varying expected read depth (25×, 100×, 200×, or 1000×). We simulated all instances by varying the value of each parameter and fixing all other parameters to default values. Briefly, we simulated each input by partitioning the SNVs into the given number of clusters and by simulating an evolutionary process where cells accumulate clusters of SNVs as well as CNAs (whole chromosome, whole arm, and focal events) which may amplify or delete overlapping SNVs. To simulate the sequencing of every SNV in each sample, we draw the total number of reads from a Poisson distribution with rate equal to the coverage and the number of variant reads from a binomial distribution, as done in previous studies ^48, 51, 53^.

We compared DeCiFer with PyClone^38^, which is the most commonly used method for direct estimation and clustering of CCFs. We ran PyClone with default parameters using two standard procedures to obtain the input: directly providing CNAs in input or providing the CCFs computed using an existing CMM-based method that accounts for subclonal CNAs^41^ (denoted as “PyClone+” in Figure 2). Further details are in Appendix B.8. We evaluated the results by computing the adjusted Rand index as well as the precision and recall across all pairs of SNVs. To calculate precision, we record the proportion of pairs that are correctly placed in the same inferred cluster, and to calculate recall we record the proportion of pairs that are in the same true cluster that were placed in the same inferred cluster.

**Figure 2:**
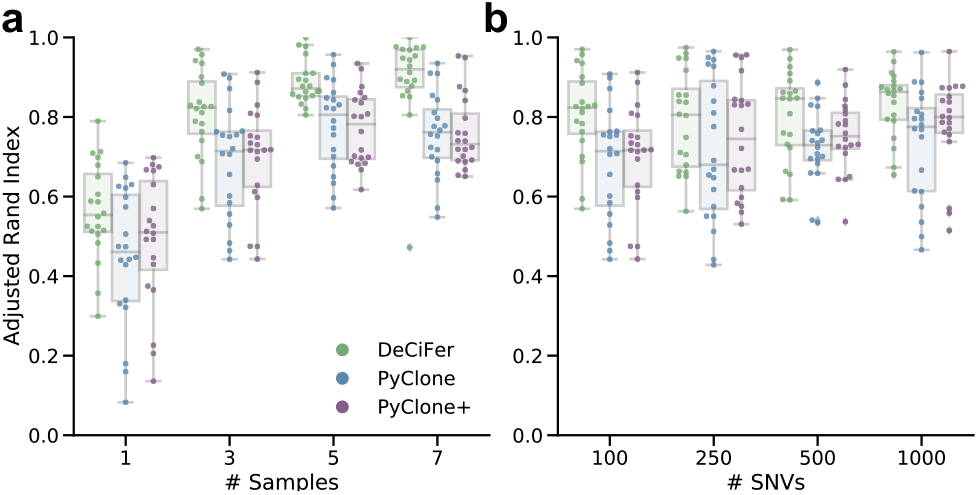
DeCiFer accurately infers clusters of SNVs on simulated data with SNV losses. **a**, The adjusted Rand index of mutation clusters computed using DeCiFer, PyClone, and PyClone+ (using CCFs computed with a CMM-based method^41^) on simulated data with varying number of samples. **b**, Same as **a** but varying number of SNVs. Each jitter-boxplot corresponds to twenty simulated data instances. The remaining parameters are fixed to default values (100 × coverage, 5 clusters, and 100 SNVs in **a** and 3 samples in **b**.)

We observe that DeCiFer consistently outperforms PyClone in terms of the ARI (Figures 2, S2). We see that performance generally increases with larger numbers of samples (Figure S2a), higher read depth (Figure S2a), and lower numbers of clusters (Figure S2b). The number of SNVs minimally affects the performance of DeCiFer and PyClone (Figure 2b). We also find that DeCiFer achieves higher clustering recall without loss of precision (Figure S3).

We also compared DeCiFer with PhyloWGS^48^, a method that infers tumor phylogenies while simultaneously clustering SNVs into clones. We ran PhyloWGS on simulated instances with 100 and 1000 SNVs. While reconstruction of a tumor phylogeny jointly from CNAs and all SNVs should in principle produce the most accurate mutation multiplicities, we found that on instances with 100 SNVs, PhyloWGS had the lowest performance (Figure S4). On instances with 1000 SNVs, we found that PhyloWGS did not converge in reasonable time (<3 days). These results suggest that phylogeny inference is challenging for large numbers of SNVs in the presence of CNAs, leading to convergence issues for approaches like PhyloWGS that attempt to simultaneously cluster SNVs and infer phylogenetic tree relating these clusters. In contrast, DeCiFer completed all instances in under 20 minutes and produced highly accurate clusters. This suggests that its approach of simultaneous selection of genotype trees for individual SNVs to maximize clustering of DCF values retains many of the advantages of full phylogenetic inference.

### 3.2 Metastatic Prostate Cancer

We analyzed SNVs and CNAs identified in whole-genome sequencing data of 49 tumor samples from 10 metastatic prostate cancer patients from Gundem et al^17^. The initial published analysis^17^ of these patients inferred CCFs for a subset of validated SNVs using the CMM assumption, clustered these SNVs into tumor clones according to their CCF values, and built a phylogenetic tree describing the evolution of these clones. We analyzed SNVs and CNAs from another published analysis of these same samples^32^, computing the DCF of each SNV using DeCiFer and computing the CCF of each SNV using the method from Dentro et al.^41^ that relies on the CMM assumption. Further details of the analysis are in Appendix B.9.

We found that the DCFs computed by DeCiFer are substantially different from the CMM CCFs for a large number of SNVs (Figure 3 and Figure S5). We summarize these differences according to two commonly used classifications of SNVs. First, we classify SNVs as *clonal* if they are inferred to be present in all cells in a tumor sample (CCF ≈ 1) or *subclonal* if they are inferred to be present only in a subpopulation (CCF ≪ 1). The clonal/subclonal classification categorizes SNVs according to their presence in cancer cells at the time of sampling. Second, we classify SNVs as *truncal* if they are inferred to occur before the most recent common ancestor of all cancer cells in the sample, and *subtrunal* otherwise. Most previous studies^16,19,21,35,38–44,47^ assume that clonal SNVs and truncal SNVs are identical. However, this assumption does not necessarily hold when SNVs are lost due to deletions in cancer cells. Using CCFs, one cannot distinguish such lost mutations from mutations that never occurred in a sample. However, the DCFs of these two cases will be different. Thus, by computing the DCF, DeCiFer has the ability to more precisely designate SNVs as truncal, especially those that were subsequently deleted and are subclonal. We classify SNVs as *truncal* if DCF ≈ 1 or *subtruncal* if DCF ≪ 1 (Figure 3a).

**Figure 3:**
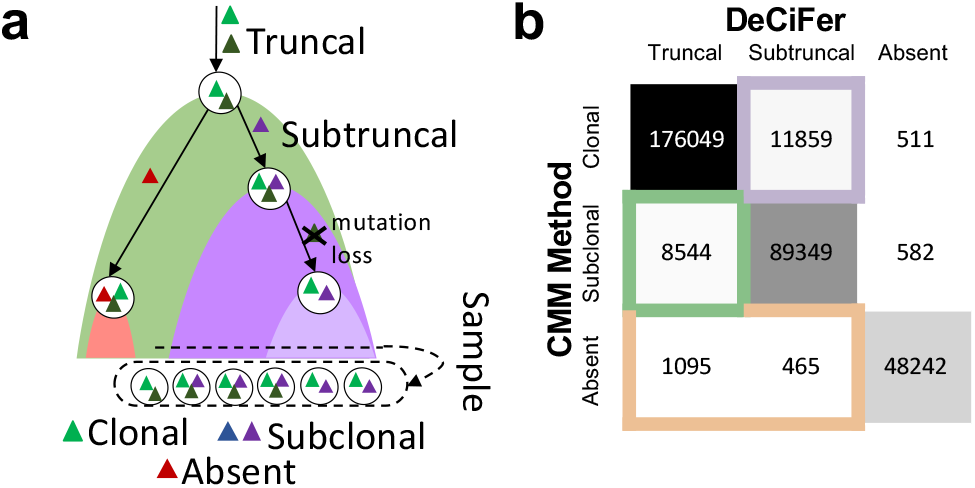
DeCiFer changes the classification of a large number of SNVs. **a**, The classification of SNVs depends on the values of their cell fractions (i.e., CCF or DCF). SNVs are classified as clonal/subclonal based on their CCF in a sample. SNVs are classified as truncal/subtruncal based on their DCF, which quantifies evolutionary history with respect to the cells in the sample. **b**, Numbers of SNVs with different classification across 49 samples from 10 prostate cancer patients.

Overall, >23000 SNVs across all samples had a change in classification (Figure 3a) when using CCFs with the CMM assumption versus DCFs inferred by DeCiFer. Multiple factors contribute to such changes. First, we found that ∼8500 SNVs across all samples that were classified as subclonal using CMM CCFs are classified as truncal by DeCiFer (Figure 3b). This difference is due to the loss of mutations by deletions. Such losses affect a moderate percentage of SNVs across all patients (5–32%, Figure S5) consistent with other recent estimates of the frequency of mutation losses^34^. We also see a large number of classification differences that cannot be explained by losses of SNVs. For example, we found that ∼12000 SNVs across all samples are classified as clonal using CMM CCFs and as subtruncal by DeCiFer (Figure 3b). These correspond to a moderate percentage of SNVs across all patients (3–40%, Figure S5). These differences are explained by choices of different mutation multiplicities (and different genotype sets r), for these SNVs. We will show below that the genotypes selected by DeCiFer often result in more parsimonious explanations for the observed data.

Another key difference in classification of SNVs is the classification of mutations as absent in a sample. SNVs without any observed variant reads in a sample (VAF = 0) are typically classified as absent from the sample. As a result, current cancer sequencing studies generally assume in downstream analyses that these SNVs were never present in the observed cancer cells in that sample or their ancestors. However, SNVs can be deleted by CNAs during tumor evolution and, when all cancer cells in a sample have been affected by such deletions, truncal SNVs may appear as absent. We find that 1, 560 SNVs across all samples classified as absent using the CMM CCFs are classified as truncal or subtruncal by DeCiFer (Figure 3b), corresponding to 0–5% of the total number of SNVs across patients (Figure S5c).

The differences between the inferred CMM CCFs and the DCFs do not simply result in different classifications of SNVs but also have a critical impact on downstream phylogenetic analysis. For example, on chromosome 5q in prostate cancer patient A17, there are two groups of 284 SNVs with different VAFs in sample A17-D (Figure 4a, top). While one group (green) is classified as clonal and the other group (brown) as subclonal based on CMM CCFs, DeCiFer classifies both groups as truncal. A previous copy-number analysis^32^ identified the presence of cancer cells with different copy numbers in the same region (Figure 4a, bottom): 61% of cancer cells have a copy-neutral loss-of-heterozygosity (LOH) (i.e., copy numbers (2, 0)), while the remaining cells are heterozygous diploid (i.e., copy numbers (1, 1)). Following the CMM assumption, the mutation multiplicity of the clonal cluster (green) is 2 in all cancer cells (Figure 4b), indicating the presence of cells with genotype (1, 1, 2). As each of these SNVs are present on both the two copies of the locus, this implies that each of the 142 SNVs occured twice (*homoplasy*), once on each homologous chromosome. While recurrent mutations, or homoplasy, may occasionally happen in tumor evolution^54^, observing homoplasy in 142 SNVs – all on the same chromosomal arm – seems *highly* unlikely.

**Figure 4:**
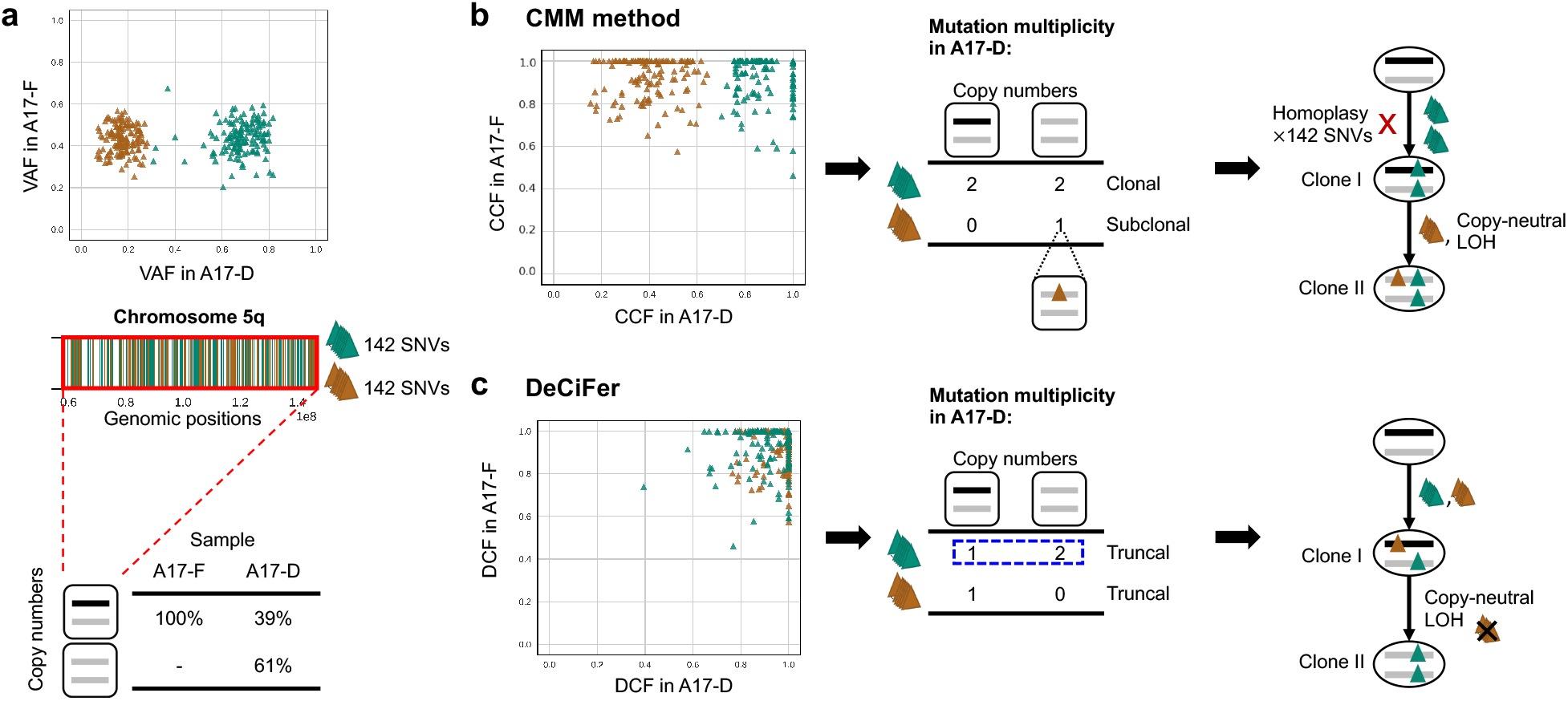
DeCiFer reclassifies subclonal mutations from CMM CCFs as truncal resulting in simpler evolutionary scenarios on prostate cancer patient A17. **a**, (Top) Two groups of 284 SNVs on chromosome 5q (brown and green) have similar VAFs in sample A17-F but different VAFs in sample A17-D, while being affected by the same CNA. (Middle) Positions of green and brown SNVs on chromosome 5q. (Bottom) Inferred copy numbers in samples: all cells in A17-F are heterozygous diploid, while 61% of cancer cells in A17-D have a copy-neutral LOH. **b**, A CMM-based method^41^ infers CCFs that separate the 284 SNVs into a clonal cluster (green) and a subclonal cluster (brown). Following the CMM assumption, the clonal SNVs (green) are inferred to have mutation multiplicity of 2 in two distinct tumor clones (clone I and II). This leads to a phylogenetic reconstruction (right) with the unrealistic conclusion that each of the 142 SNVs occurred twice independently on both homologous chromosomes (i.e., 142 homoplasy events). **c**, DeCiFer infers DCF≈1 for all 284 SNVs by identifying different mutation multiplicities (blue dashed box) for SNVs in the green cluster across the two clones (I and II) and by identifying loss of brown SNVs in subset of tumor cells. Thus all SNVs in the green and brown cluster are truncal. This results in a realistic tumor phylogeny where the mutation multiplicities are consistent with the observed copy-neutral LOH.

On the same patient A17, DeCiFer infers DCFs that result in a much more realistic mutation multiplicies and phylogenetic reconstruction. In particular, DeCiFer infers that the two groups of SNVs (green and brown in Figure 4a) are part of the same truncal cluster (i.e., DCF≈ 1) but have different mutation multiplicities: the SNVs in the first group (green) have a multiplicity of 1 in one clone and a multiplicity of 2 in the other, a scenario that is not allowed under the CMM assumption. DeCiFer’s results are consistent with a realistic evolutionary scenario where the two groups of SNVs occurred on different alleles of the chromosome: the first group (green) was amplified during the copy-neutral LOH event, while the second group (brown) was lost. Further supporting this reconstruction is the observation that the two groups of SNVs are randomly distributed over chromosome 5q (Figure 4a), indicating that the differences in VAF between the green and brown SNVs are not due to an error in the copy numbers for one group. Thus, DeCiFer’s classification of the SNVs in the brown group as truncal, but lost in a subpopulation of cancer cells results in a simpler and more realistic reconstruction of tumor evolution compared to the classification of these SNVs as subclonal according to their CMM CCFs. Notably, on another patient (A-24), we observe the opposite difference in classification: SNVs classified as clonal by the CMM CCFs are classified as subtruncal by DeCiFer resulting similarly in a simpler reconstruction of tumor evolution (Section C.2 and Figure S6).

On prostate cancer patient A12, we see a example of mutations that are classified as absent using CMM CCFs but classified as truncal by DeCiFer. Specifically, Chromosome 6q contains 86 SNVs that are split into two groups with different VAFs across three samples (Figure 5a, top): the first group (green) has VAFs between 0.4–0.8 in all samples, while the second group (magenta) has VAFs between 0.1–0.4 in samples A12-C and A12-D but VAFs ≈ 0 in the remaining sample A12-A. These two groups of SNVs have different CMM CCFs across samples and result in the inference of three distinct tumor clones with a complicated evolution characterized by recurrent mutation (homoplasy) of 58 SNVs in the magenta cluster (Figure 5b). However, a previous copy-number analysis^32^ revealed the presence of a copy-neutral LOH on chromosome 6q in these samples. The proportions of cancer cells that have the LOH closely follow the VAFs of the SNVs in the magenta group of SNVs (Figure 5a, bottom): 100%, 66%, and 0% of cancer cells have a copy-neutral LOH in A12-A, A12-C, and A12-D, respectively. DeCiFer infers that all the 86 SNVs in the green and magenta groups are truncal with DCFs ≈ 1, and that the magenta group of SNVs were lost during the copy-neutral LOH event (Figure 5c). Indeed, in sample A12-A, where all cancer cells have the copy-neutral LOH, these SNVs have a VAF= 0. Note that the phyloCCF correction introduced in Jamal-Hanjani et al.^13^ to address issues of mutation loss would not have identified the magenta SNVs as truncal. Since the phyloCCF is applied on each sample independently, it cannot distinguish between mutation losses and absences in a given sample, and thus is only applied to SNVs with VAF > 0. Thus DeCiFer yields a more parsimonious reconstruction of tumor evolution than obtained with CMM CCFs, with fewer tumor clones and no massive homoplasy.

**Figure 5:**
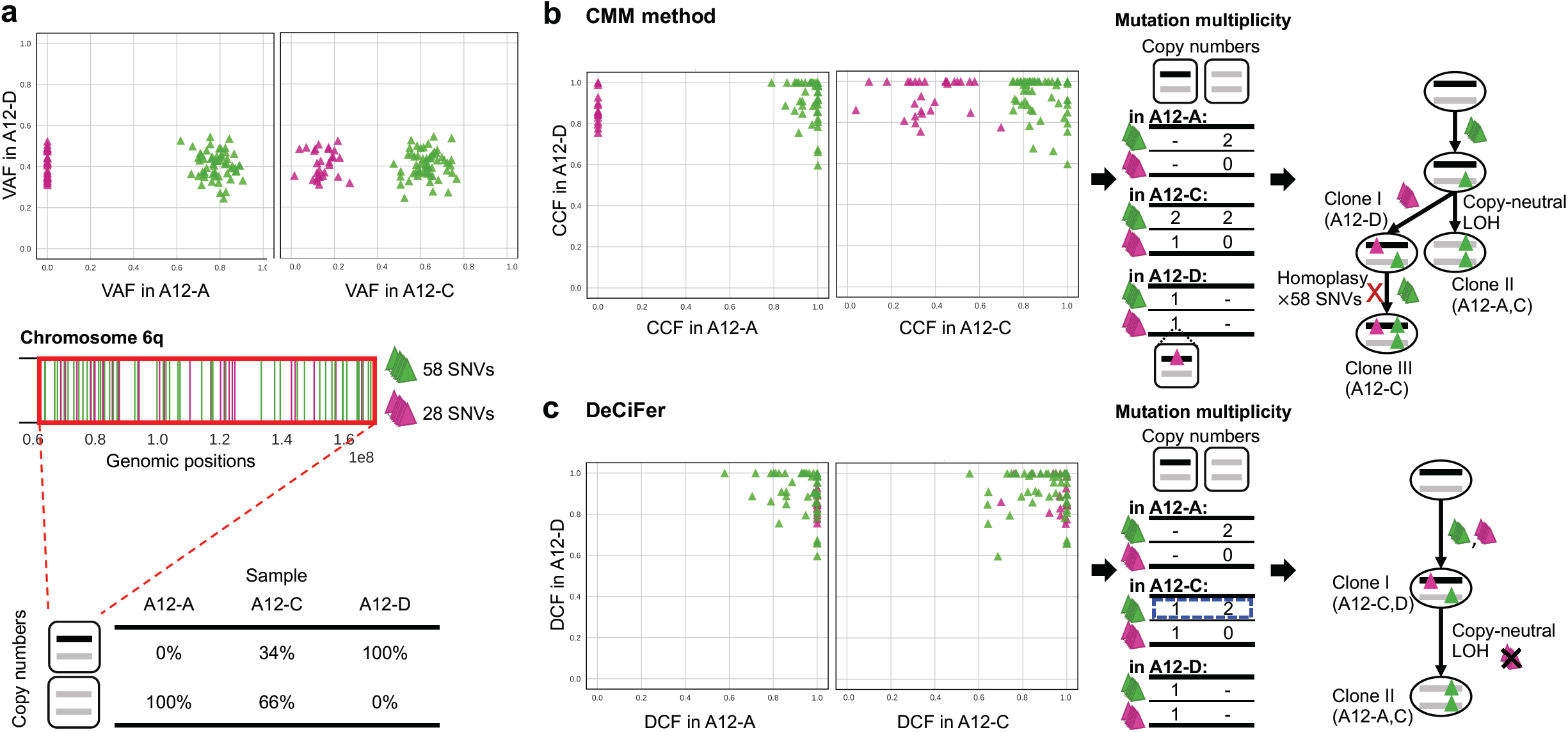
DeCiFer accurately identifies losses of SNVs in prostate cancer patient A12. **a**, Two groups of 87 SNVs on chromosome 6q (green and magenta) have different VAFs in samples A12-A and A12-C, with one group (magenta) having VAF =0 in A12-A. (Middle) Positions of green and magenta SNVs on chromosome 6q. (Bottom): Inferred copy numbers in samples: sample A12-A only contains cancer cells with a copy-neutral LOH, A12-D only contains heterozygous diploid cells, and A12-C contains a mixture of both. **b**, A CMM-based method^41^ infers CCFs that separate the SNVs into a clonal cluster in all samples (green) and another cluster (magenta), which is clonal in A12-D, absent in A12-A, and subclonal in A12-C. Following the CMM assumption, the mutation multiplicities in each sample indicate the presence of three distinct tumor clones (labeled I, II, and III). This leads to a phylogenetic reconstruction (right) with the unrealistic conclusion that 58 SNVs occurred twice independently on both homologous chromosomes in Clone III (58 homoplasy events). **c**, DeCiFer infers DCF ≈ 1 for all SNVs by identifying different mutation multiplicities (blue dashed box) for SNVs in green cluster in sample A12-C and by identifying the loss of 28 magenta SNVs in a subset of tumor cells due to a copy-neutral LOH. Thus, all SNVs in green and magenta clusters are truncal. Notably, these 28 SNVs have DCF ≈ 1 also in A12-A even though their VAF = 0; this is because all cancer cells have a copy-neutral LOH in sample A12-A. The inferred losses of these 28 SNVs are well supported by the fact that the proportion of cancer cells with the copy-neutral LOH (second row of table in (**a**)) closely follows their variations of VAF across samples (VAFs in (**a**)).

## 4 Discussion

The cancer cell fraction (CCF) is the cornerstone of tumor heterogeneity and evolution analysis using single-nucleotide variants (SNV). However, we demonstrated that current approaches to estimate CCFs suffer from major limitations and these limitations have striking consequences on real data. First, nearly all existing methods to estimate CCFs are based on the Constant Mutation Multiplicity (CMM) assumption that is violated in many tumors. Second, the CCF is not the correct quantity to group mutations for phylogenetic analysis in the case where SNV losses occur due to CNAs, a case that is common in solid tumors. In this work, we address these limitations by: (i) introducing the Single-Split Copy Number (SSCN) assumption, a more realistic alternative to the CMM assumption; (ii) defining the descendant cell fraction (DCF), a generalization of the CCF that accounts for SNV losses; (iii) developing DeCiFer, an algorithm that simultaneously estimates DCFs (or CCFs) of individual SNVs and clusters SNVs into a small number of groups according to these DCFs (or CCFs) across multiple tumor samples. We show that DeCiFer improves the identification of SNVs that co-occurred in the same tumor clone on both simulated and real cancer data, yielding more realistic reconstructions of tumor evolution compared to earlier approaches based on CCFs inferred using the CMM assumption. DCF clusters account for mutation losses and differences in copy number, and thus can be used as input to standard tumor phylogeny methods^55–59^. This will enable phylogeny inference for realistic sized problems containing thousands of SNVs whose copy numbers may differ within and across tumor samples. DeCiFer can also be run with CCFs instead of DCFs, which may be preferable for certain applications such as neoantigen prediction^60^ where identifying the clonal status of mutations is important.

While DeCiFer enables us to overcome some major limitations of previous studies, there are several venues for future improvements. First, we assume that the given copy numbers and proportions are exact. However, methods that infer copy numbers and proportions from bulk DNA sequencing data are subject to errors and may miss CNAs that are small or present at low proportion in a sample. This uncertainty could be incorporated into the DeCiFer model. Second, further improvements in SNV clustering are possible, such as better modeling of the tail of low prevalence SNVs that are expected due to neutral evolution^47^. Third, breakpoints of structural variants could also be analyzed by DeCiFer since the prevalence of these mutations is proportional to read counts^61^. Finally, the genotype trees selected by DeCiFer for each SNV could be combined into tumor phylogenies, perhaps using consenus tree methods. DeCiFer provides a robust framework to decipher tumor heterogeneity in the presence of copy number aberrations providing a tool to improving understanding of tumor development and evolution.

## References

[1] Nowell, P. C. The clonal evolutiJon of tumor cell populations. Science 194, 23–8 (1976).

[2] Burrell, R. A., McGranahan, N., Bartek, J. & Swanton, C. The causes and consequences of genetic heterogeneity in cancer evolution. Nature 501, 338–345 (2013).

[3] Andor, N. et al. Pan-cancer analysis of the extent and consequences of intratumor heterogeneity. Nature Medicine 22, 105–113 (2015).

[4] McGranahan, N. & Swanton, C. Clonal Heterogeneity and Tumor Evolution: Past, Present, and the Future. Cell 168, 613–628 (2017).

[5] Navin, N. E. The first five years of single-cell cancer genomics and beyond. Genome research 25, 1499–1507 (2015).

[6] Gawad, C., Koh, W. & Quake, S. R. Single-cell genome sequencing: current state of the science. Nature Reviews Genetics 17, 175 (2016).

[7] Kim, C. et al. Chemoresistance evolution in triple-negative breast cancer delineated by single-cell sequencing. Cell 173, 879–893 (2018).

[8] Gao, R. et al. Punctuated copy number evolution and clonal stasis in triple-negative breast cancer. Nature genetics 48, 1119 (2016).

[9] Laks, E. et al. Clonal decomposition and DNA replication states defined by scaled single-cell genome sequencing. Cell 179, 1207–1221 (2019).

[10] Myers, M. A., Zaccaria, S. & Raphael, B. J. Identifying tumor clones in sparse single-cell mutation data. Bioinformatics 36, i186–i193 (2020).

[11] Zaccaria, S. & Raphael, B. J. Characterizing allele-and haplotype-specific copy numbers in single cells with CHISEL. Nature Biotechnology 1–8 (2020).

[12] The, I., of Whole, T. P.-C. A., Consortium, G. et al. Pan-cancer analysis of whole genomes. Nature 578, 82 (2020).

[13] Jamal-Hanjani, M. et al. Tracking the Evolution of Non-Small-Cell Lung Cancer. New England Journal of Medicine 376, 2109–2121 (2017).

[14] Dewey, F. E. et al. Clinical interpretation and implications of whole-genome sequencing. Jama 311, 1035–1045 (2014).

[15] Rajput, A., Bocklage, T., Greenbaum, A., Lee, J.-H. & Ness, S. A. Mutant-allele tumor heterogeneity scores correlate with risk of metastases in colon cancer. Clinical colorectal cancer 16, e165–e170 (2017).

[16] Dentro, S. C. et al. Characterizing genetic intra-tumor heterogeneity across 2,658 human cancer genomes. bioRxiv (2020).

[17] Gundem, G. et al. The evolutionary history of lethal metastatic prostate cancer. Nature 520, 353–357 (2015).

[18] Brastianos, P. K. et al. Genomic Characterization of Brain Metastases Reveals Branched Evolution and Potential Therapeutic Targets. Cancer Discovery 5, 1164–1177 (2015).

[19] Cresswell, G. D. et al. Mapping the breast cancer metastatic cascade onto ctdna using genetic and epigenetic clonal tracking. Nature communications 11, 1–12 (2020).

[20] Williams, M. J., Werner, B., Barnes, C. P., Graham, T. A. & Sottoriva, A. Identification of neutral tumor evolution across cancer types. Nature Genetics 48, 238–244 (2016).

[21] Lakatos, E. et al. Evolutionary dynamics of neoantigens in growing tumors. Nature Genetics 1–10 (2020).

[22] Bozic, I. et al. Accumulation of driver and passenger mutations during tumor progression. Proceedings of the National Academy of Sciences 107, 18545–18550 (2010).

[23] Rubanova, Y. et al. Reconstructing evolutionary trajectories of mutation signature activities in cancer using tracksig. Nature communications 11, 1–12 (2020).

[24] Harrigan, C. F., Rubanova, Y., Morris, Q. & Selega, A. Tracksigfreq: subclonal reconstructions based on mutation signatures and allele frequencies. In Pacific Symposium on Biocomputing. Pacific Symposium on Biocomputing, vol. 25, 238 (World Scientific, 2020).

[25] Christensen, S., Leiserson, M. D. & El-Kebir, M. Physigs: Phylogenetic inference of mutational signature dynamics. In Pacific Symposium on Biocomputing. Pacific Symposium on Biocomputing, vol. 25, 226–237 (World Scientific, 2020).

[26] Van Loo, P. et al. Allele-specific copy number analysis of tumors. Proceedings of the National Academy of Sciences 107, 16910–16915 (2010).

[27] Carter, S. L. et al. Absolute quantification of somatic DNA alterations in human cancer. Nature Biotechnology 30, 413–421 (2012).

[28] Nik-Zainal, S. et al. The life history of 21 breast cancers. Cell 149, 994–1007 (2012).

[29] Oesper, L., Mahmoody, A. & Raphael, B. J. THetA: inferring intra-tumor heterogeneity from high-throughput DNA sequencing data. Genome Biol 14, R80 (2013).

[30] Oesper, L., Satas, G. & Raphael, B. J. Quantifying tumor heterogeneity in whole-genome and whole-exome sequencing data. Bioinformatics 30, 3532–40 (2014).

[31] Fischer, A., Vazquez-Garcia, I., Illingworth, C. J. R. & Mustonen, V. High-definition reconstruction of clonal composition in cancer. Cell Reports 7, 1740–1752 (2014).

[32] Zaccaria, S. & Raphael, B. J. Accurate quantification of copy-number aberrations and whole-genome duplications in multi-sample tumor sequencing data. Nature Communications 11, 1–13 (2020).

[33] Zack, T. I. et al. Pan-cancer patterns of somatic copy number alteration. Nature genetics 45, 1134–1140 (2013).

[34] McPherson, A. et al. Divergent modes of clonal spread and intraperitoneal mixing in high-grade serous ovarian cancer. Nature Genetics (2016).

[35] Gerstung, M. et al. The evolutionary history of 2,658 cancers. Nature 578, 122–128 (2020).

[36] Watkins, T. B. et al. Pervasive chromosomal instability and karyotype order in tumour evolution. Nature 587, 126–132 (2020).

[37] Weaver, B. A. & Cleveland, D. W. Does aneuploidy cause cancer? Current opinion in cell biology 18, 658–667 (2006).

[38] Roth, A. et al. PyClone: statistical inference of clonal population structure in cancer. Nature methods 11, 396–398 (2014).

[39] Miller, C. A. et al. SciClone: inferring clonal architecture and tracking the spatial and temporal patterns of tumor evolution. PLoS Comput Biol 10, e1003665 (2014).

[40] McGranahan, N. et al. Clonal status of actionable driver events and the timing of mutational processes in cancer evolution. Science Translational Medicine 7, 283ra54–283ra54 (2015).

[41] Dentro, S. C., Wedge, D. C. & Van Loo, P. Principles of Reconstructing the Subclonal Architecture of Cancers. Cold Spring Harbor Perspectives in Medicine 7, a026625 (2017).

[42] Yuan, K. et al. Ccube: a fast and robust method for estimating cancer cell fractions. bioRxiv 484402 (2018).

[43] Cun, Y., Yang, T.-P., Achter, V., Lang, U. & Peifer, M. Copy-number analysis and inference of subclonal populations in cancer genomes using sclust. Nature protocols 13, 1488 (2018).

[44] Xiao, Y. et al. Fastclone is a probabilistic tool for deconvoluting tumor heterogeneity in bulk-sequencing samples. Nature Communications 11, 1–11 (2020).

[45] Tarabichi, M. et al. A practical guide to cancer subclonal reconstruction from DNA sequencing. Nature Methods (2021). URL http://www.nature.com/articles/s41592-020-01013-2.

[46] Yuan, K., Sakoparnig, T., Markowetz, F. & Beerenwinkel, N. BitPhylogeny: a probabilistic framework for reconstructing intra-tumor phylogenies. Genome biology 16, 1 (2015).

[47] Caravagna, G. et al. Subclonal reconstruction of tumors by using machine learning and population genetics. Nature Genetics 52, 898–907 (2020).

[48] Deshwar, A. G. et al. PhyloWGS: Reconstructing subclonal composition and evolution from wholegenome sequencing of tumors. Genome Biology 16, 35 (2015).

[49] El-Kebir, M., Satas, G., Oesper, L. & Raphael, B. J. Inferring the Mutational History of a Tumor Using Multi-state Perfect Phylogeny Mixtures. Cell Systems 3, 43–53 (2016).

[50] Jiang, Y., Qiu, Y., Minn, A. J. & Zhang, N. R. Assessing intratumor heterogeneity and tracking longitudinal and spatial clonal evolutionary history by next-generation sequencing. Proceedings of the National Academy of Sciences of the United States of America 113, E5528–37 (2016).

[51] Satas, G., Zaccaria, S., Mon, G. & Raphael, B. J. Scarlet: Single-cell tumor phylogeny inference with copy-number constrained mutation losses. Cell Systems 10, 323–332 (2020).

[52] El-Kebir, M. SPhyR: tumor phylogeny estimation from single-cell sequencing data under loss and error. Bioinformatics 34, i671–i679 (2018). URL https://academic.oup.com/bioinformatics/article/34/17/i671/5093218.

[53] Satas, G. & Raphael, B. J. Tumor phylogeny inference using tree-constrained importance sampling. Bioinformatics 33, i152–i160 (2017).

[54] Kuipers, J., Jahn, K., Raphael, B. J. & Beerenwinkel, N. Single-cell sequencing data reveal widespread recurrence and loss of mutational hits in the life histories of tumors. Genome research 27, 1885–1894 (2017).

[55] Popic, V. et al. Fast and scalable inference of multi-sample cancer lineages. Genome biology 16, 91 (2015).

[56] Qiao, Y. et al. SubcloneSeeker: a computational framework for reconstructing tumor clone structure for cancer variant interpretation and prioritization. Genome biology 15,443 (2014).

[57] El-Kebir, M., Oesper, L., Acheson-Field, H. & Raphael, B. J. Reconstruction of clonal trees and tumor composition from multi-sample sequencing data. Bioinformatics 31, i62–i70 (2015).

[58] Malikic, S., McPherson, A. W., Donmez, N. & Sahinalp, C. S. Clonality inference in multiple tumor samples using phylogeny. Bioinformatics 31, 1349–1356 (2015).

[59] Husic, E. et al. Mipup: minimum perfect unmixed phylogenies for multi-sampled tumors via branchings and ilp. Bioinformatics 35, 769–777 (2019).

[60] Cai, W. et al. Mhc class ii restricted neoantigen peptides predicted by clonal mutation analysis in lung adenocarcinoma patients: implications on prognostic immunological biomarker and vaccine design. BMC genomics 19, 1–9 (2018).

[61] Cmero, M. et al. Inferring structural variant cancer cell fraction. Nature communications 11, 1–15 (2020).

[62] Virtanen, P. et al. SciPy 1.0: Fundamental Algorithms for Scientific Computing in Python. Nature Methods 17, 261–272 (2020).

[63] Thorndike, R. L. Who belongs in the family? Psychometrika 18, 267–276 (1953).

[64] Zhang, Y., Maiidziuk, J., Quek, C. H. & Goh, B. W. Curvature-based method for determining the number of clusters. Information Sciences 415, 414–428 (2017).

[65] Koboldt, D. C. et al. VarScan 2: somatic mutation and copy number alteration discovery in cancer by exome sequencing. Genome Research 22, 568–576 (2012).

