## Appendix for "DeCiFering the Elusive Cancer Cell Fraction in Tumor Heterogeneity and Evolution"

### A Supplementary Figures

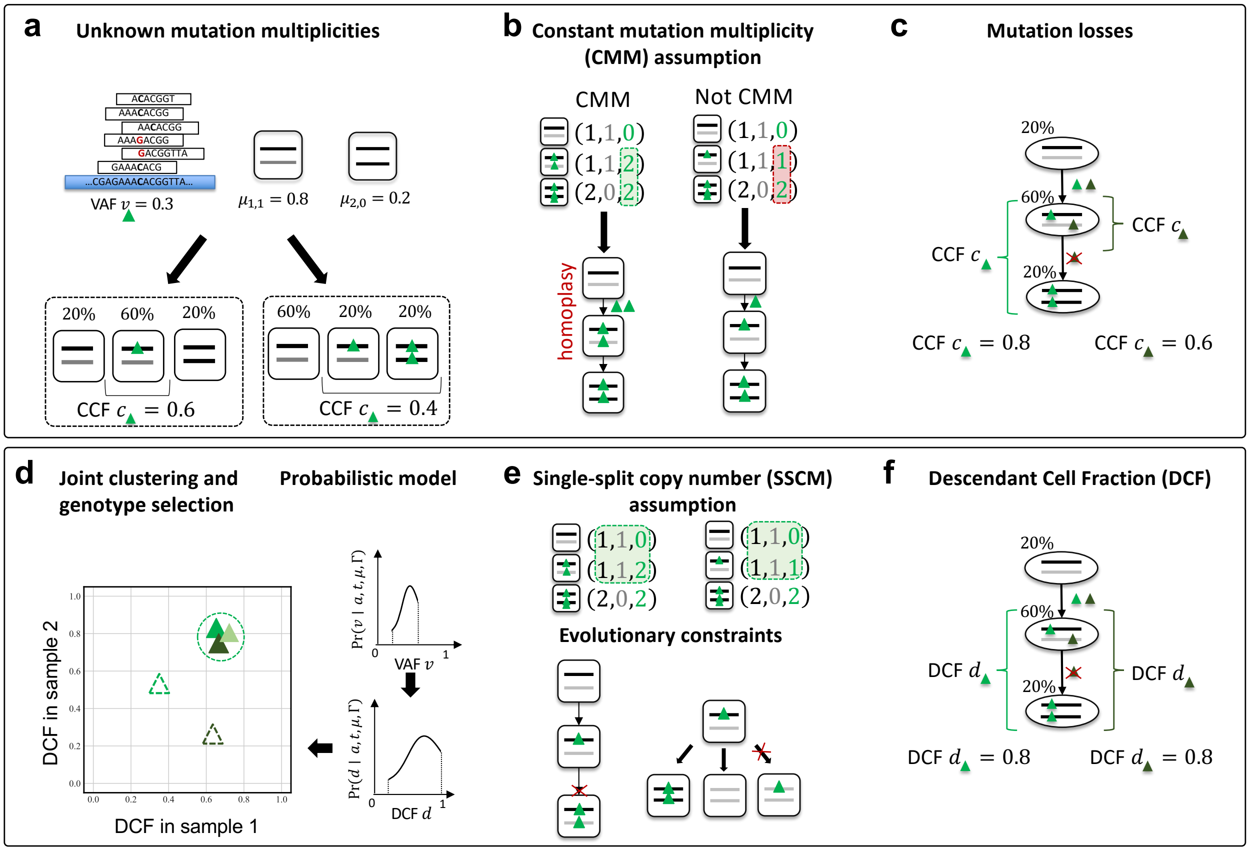

**Figure S1: Limitations of the cancer cell fraction and the contributions of DeCiFer.** **a**, Our observations from sequencing data correspond to the total and variant number of sequencing reads from the genomic locus of each somatic SNV (green triangle), which we use to obtain an estimate of VAF. Moreover, existing methods also enable us to identify the whole-genome copy numbers and proportions of the different cells. However, these values do not directly yield the cancer cell fraction, as the number of mutations in each clone, i.e., the mutation multiplicities, are unknown. As such, there may be many possible CCF values for the same observed VAF and copy-number states. **b**, Most existing approaches to estimate CCF rely on the constant mutation multiplicity assumption, which states that all clones that have the mutation have the same number of copies of the mutation. However, this assumption may admit sets of genotypes (left) that are evolutionary unlikely. Moreover, it may also exclude sets of genotypes (right) that are likely. **c**, In addition, CCF does not directly account for mutation losses. Groups of SNVs that co-occur on the same branch of a phylogeny may have different CCFs due to mutation loss. **d**, DeCiFer addresses the problem of unknown mutation multiplicities, by assuming that mutations derive from a set of clusters and simultaneously selecting mutation multiplicites (i.e., genotype selection) and clustering, using a probabilistic model of sequencing data. **e**, Instead of the CMM assumption, we make the single-split copy-number (SSCM) assumption. In addition, we use a set of evolutionary constraints to restrict sets of genotypes, e.g., each mutation occurs at most once and mutation multipliciy changes only due to copy-number aberrations. **f**, To account for possible mutation loss, we introduce the Descendant Cell Fraction (DCF). The DCF of an SNV is defined as the proportion of cells in a sample that are descendant to the cell that acquired the SNV.

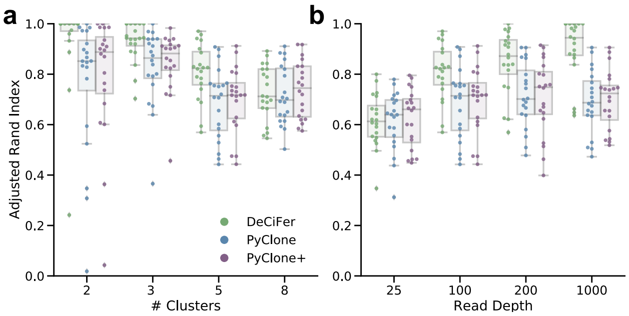

**Figure S2: DeCiFer accurately infers clusters of SNVs on simulated data with SNV losses.** Plots show the adjusted Rand index for DeCiFer, PyClone, and PyClone+ (using CCFs computed with the method outlined by Dentro et al.[41]) results on simulated data. Each boxplot corresponds to twenty simulated data instances. Panels **a** and **b** respectively show the results of varying the number of clusters $k$ and the expected read depth $c$. The remaining parameters are fixed to default values of $m = 3$, $k = 5$, $n = 100$ and $c = 100$.

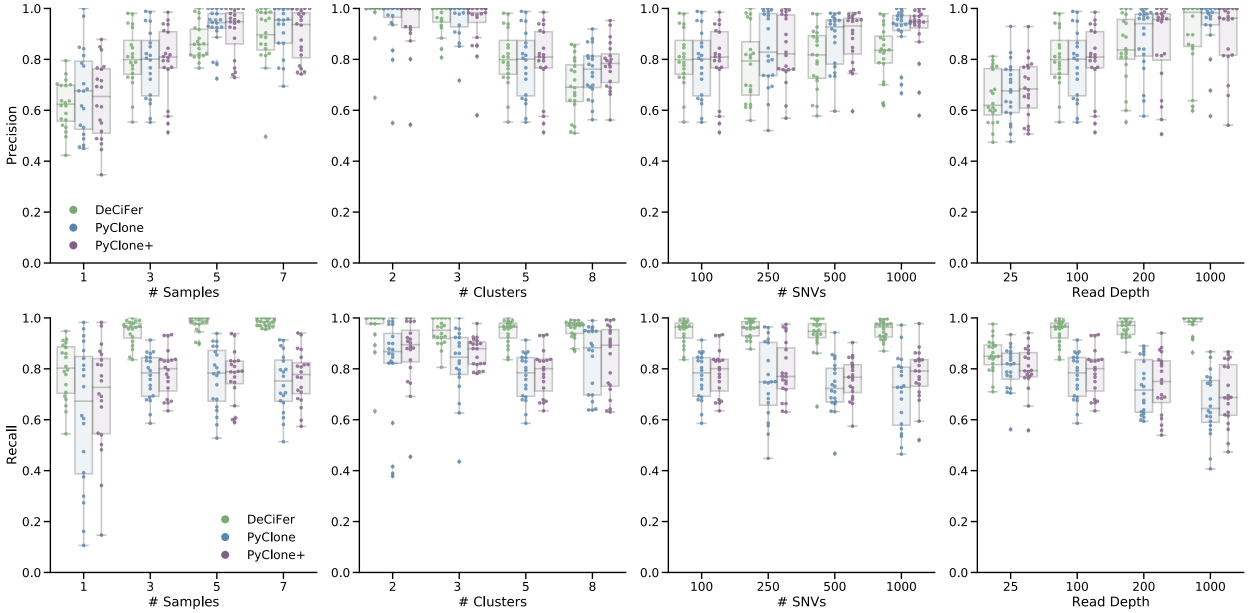

**Figure S3: DeCiFer achieves high recall for clusters of SNVs on simulated data with SNV losses.** Plots show precision and recall for DeCiFer, PyClone, and PyClone+ (using CCFs computed with the method outlined by Dentro et al.[41]) results on simulated data.

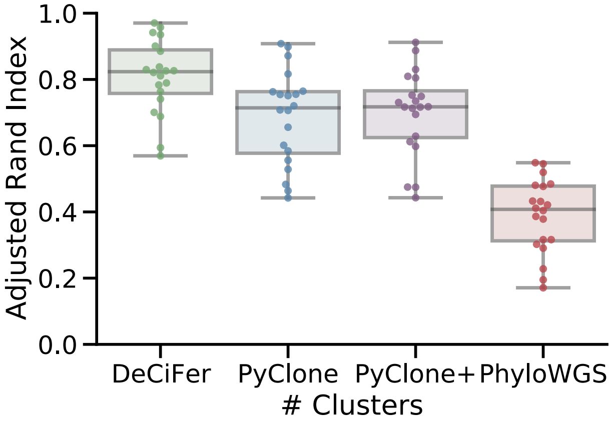

**Figure S4: DeCiFer outperforms both CMM and phylogenetic approaches for inferring SNV clusters on simulated data with SNV losses**. Plots show ARI for DeCiFer, PyClone, and PyClone+ (using CCFs computed with the method outlined by Dentro et al.[41]) and PhyloWGS results on simulated data. Each boxplot corresponds to 20 simulated data instances, with parameters $m = 3$, $k = 5$, $n = 100$ and $c = 100$

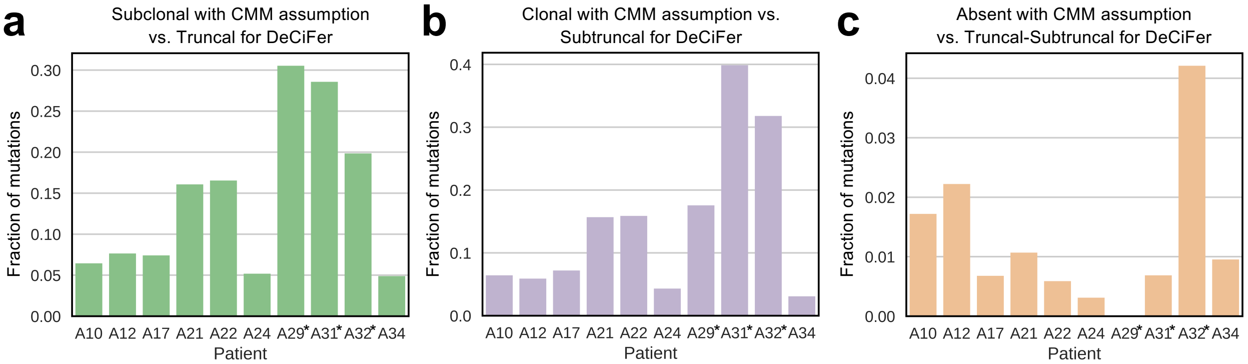

**Figure S5: DeCiFer identifies a substantial different number of truncal and subtruncal SNVs.** The classification of somatic SNVs depends on the the values of their cell fractions (i.e., CCF or DCF) across the samples of 10 prostate cancer patient, including 3 patients in which the occurrence of a WGD has been identified in previous copy-number analysis[32] (starred). **b**, Number of SNVs classified as truncal by DeCiFer (DCF$\geq$ 0.9) and as subtruncal using the CMM assumption (CCF$<$ 0.9) in at least one samples of every prostate cancer patient. **c**, Number of SNVs classified as subtruncal by DeCiFer (DCF$<$ 0.9) and as truncal using the CMM assumption (CCF$<$ 0.9) in at least one samples of every prostate cancer patient. **d**, Number of SNVs classified as either truncal or subtruncal by DeCiFer (DCF$\geq$ 0.1) and as absent using the CMM assumption (CCF$<$ 0.1) in at least one samples of every prostate cancer patient.

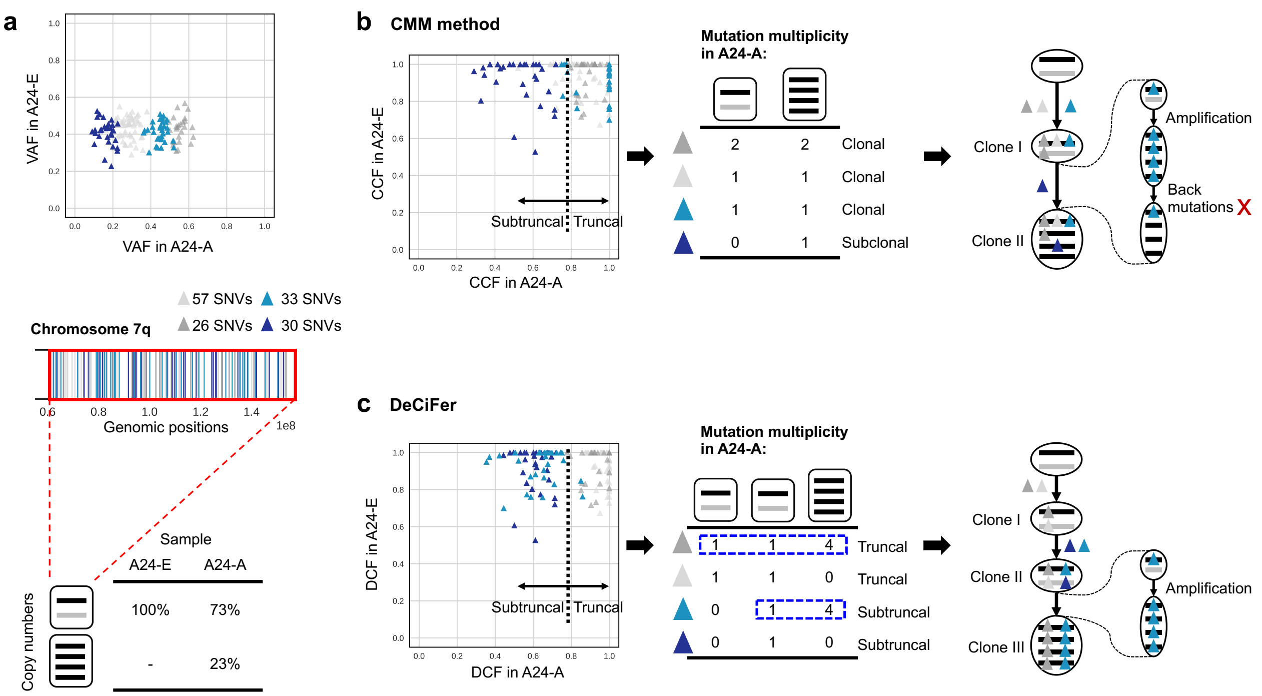

**Figure S6: DeCiFer accurately identifies subtruncal SNVs that are classified as truncal by the CMM assumption with unrealistic evolutionary scenarios.** **a**, 167 somatic SNVs are located on chromosome 7 of prostate cancer patient A24. **b**, Four clusters of these SNVs have the same values of VAF in sample A24-E but different values in A24-A. **c**, Different cells have different copy numbers in the two samples for the genomic region harboring these SNVs: while all cells in A24-E are diploid in this region (i.e., copy numbers $(1, 1)$), the 20% of cells in A24-A have a LOH with an amplification (i.e., copy numbers $(4, 0)$). **d**, An existing method[41] using the CMM assumption infers that three clusters (cyan, and light and dark grey) of SNVs in (**b**) have CCF$\approx$1 in both samples, corresponding to truncal SNVs. Instead, the method infers that the remaining cluster (dark blue) is subclonal in A24-A, which thus includes two tumor clones (clone I and II). Based on the values of VAF and the CMM assumption, the method infers that each of the three clusters of clonal SNVs have the same SNV multiplicty: two of these clusters (light grey and cyan) have SNV multiplicty of 1 and the other cluster (dark grey) has an SNV multiplicty of 2. Since all the SNVs are affected by an amplification of the retained allele and since SNVs are also amplified with the corresponding allele, this result implies the unrealistic occurrence of back mutations for all the clusters; for example, three back mutations must have occurred for one of these clusters (cyan). **e**, DeCiFer infers that two (light and dark grey) of the clusters in (**b**) have DCF$\approx$1 and correspond to truncal SNVs, while the other two remaining clusters (ligth and dark blue) have DCF$\approx$0.6 and correspond to subtruncal SNVs. Specifically, DeCiFer obtains this result by identifying the loss of two clusters (ligth gray and dark blue) located on the same allele (black) and the presence of different mutation multiplicities in different clones (clones II and III) for the other two remaining clusters (dark grey and ligth blue), resulting in a realistic tumor phylogeny.

### B Supplementary Methods

#### B.1 Derivation of Theorem 1

In this section, we derive in detail Main Text Theorem 1, which describes the relationship between CCF $c$ and VAF $v$ for an SSCN genotype set $\Gamma^*$. We will begin by restating several definitions provided in the main text. Given the tumor purity $\rho$, genotype set $\Gamma$ and genotype proportions $\mathbf{g}$ uniquely determine the CCF $c$.

**Definition 1**

The *cancer cell fraction* (CCF) $c$ is defined in terms of a genotype set $\Gamma$ and genotype proportions $\mathbf{g}$ and tumor purity $\rho$ as

$$c = \frac{1}{\rho} \sum_{(x,y,m) \in \Gamma_{\text{CCF}}} g_{(x,y,m)}, \tag{S1}$$

where $\Gamma_{\text{CCF}}$ is the subset of genotypes $(x, y, m) \in \Gamma$ where $m \geq 1$.

The genotype set $\Gamma$ and genotype proportions $\mathbf{g} = [g_{(x,y,m)}]$ uniquely determine the VAF $v$ as follows.

**Definition 2**

The *variant allele frequency* (VAF) $v$ is defined in terms of a genotype set $\Gamma$ and genotype proportions $\mathbf{g}$ as

$$v = \frac{1}{F} \sum_{(x,y,m) \in \Gamma} m \cdot g_{(x,y,m)} \tag{S2}$$

where

$$F = \sum_{(x,y,m) \in \Gamma} (x + y) \cdot g_{(x,y,m)} \tag{S3}$$

is the fractional copy number of the locus as defined by $\Gamma$ and $\mathbf{g}$.

The genotype set $\Gamma$ and genotype proportions $\mathbf{g} = [g_{(x,y,m)}]$ uniquely determine the copy-number proportions $\boldsymbol{\mu}$ as follows.

**Definition 3**

The *copy-number proportions* $\boldsymbol{\mu}$ is defined in terms of a genotype set $\Gamma$ and genotype proportions $\mathbf{g}$ as

$$\mu_{(x,y)} = \frac{1}{F} \sum_{(x,y,m) \in \Gamma} m \cdot g_{(x,y,m)} \quad \text{(S4)}$$

for all copy-number states $(x, y)$.

We define consistency between the VAF $v$, copy-number proportions $\boldsymbol{\mu}$, genotype sets $\Gamma$ and the underlying genotype states $\mathbf{g}$ in terms of the above equations.

**Definition 4**

Genotype set and proportions $(\Gamma, \mathbf{g})$ are *consistent* with VAF $v$ and copy-number proportions $\boldsymbol{\mu}$ provided that Eqs. (S2) and (S4) are satisfied.

**Definition 5**

Genotype set $\Gamma$ is *consistent* with VAF $v$ and copy-number proportions $\boldsymbol{\mu}$ provided that there exist genotype proportions $\mathbf{g}$ such that Eqs. (S2) and (S4) are satisfied.

In the main text, we introduce the Single Split Copy Number assumption, which follows from simple evolutionary models on SNVs and CNAs: the Dollo model for SNVs, and infinite alleles model for allele-specific copy numbers.

**Assumption**

**Single Split Copy Number (SSCN).** At every SNV locus, there is exactly one copy-number state $(x^*, y^*)$ with two distinct genotypes $(x^*, y^*, 0)$ and $(x^*, y^*, m^*)$.

We make the claim for SSCN genotype sets $\Gamma^*$ that provided consistent genotype proportions $\mathbf{g}$ exist, they are unique. Specifically they take the following form.

**Lemma 1**

Given VAF $v$, copy-number proportions $\boldsymbol{\mu}$, and an SSCN genotype set $\Gamma^*$, if genotype proportions $\mathbf{g}$ exist such that $(\Gamma^*, \mathbf{g})$ are consistent with VAF $v$ and copy-number proportions $\boldsymbol{\mu}$, they are uniquely determined as

$$g_{(x,y,m)} = \begin{cases} \mu_{(x,y)}, & \text{if } (x,y) \neq (x^*,y^*), \\ (1-\lambda)\cdot\mu_{(x^*,y^*)}, & \text{if } (x,y) = (x^*,y^*) \text{ and } m = 0, \\ \lambda\cdot\mu_{(x^*,y^*)}, & \text{if } (x,y) = (x^*,y^*) \text{ and } m = m^*, \end{cases} \tag{S5}$$

where

$$\lambda = \frac{1}{m^*\mu_{(x^*,y^*)}}\left[vF - \sum_{\substack{(x,y,m)\in\Gamma^* \\ (x,y)\neq(x^*,y^*)}} m\cdot\mu_{(x,y)}\right]. \tag{S6}$$

**Proof**

This follows from the definition of consistency and Eqs. (S2) and (S4). Consider the case where $(x,y) \neq (x^*,y^*)$. By the SSCN assumption, there exists a unique single $(x',y',m) \in \Gamma^*$ with $(x,y) = (x',y')$. Thus, to satisfy Eq. (S4) we have that $g_{(x,y,m)} = \mu_{(x,y)}$.

Next, consider the case where $(x,y) = (x^*,y^*)$. Thus, we have that $\mu_{(x^*,y^*)} = g_{(x^*,y^*,0)} + g_{(x^*,y^*,m^*)}$, or equivalently $g_{(x^*,y^*,0)} = (1-\lambda)\mu_{(x^*,y^*)}$ and $g_{(x^*,y^*,m^*)} = \lambda\mu_{(x^*,y^*)}$ for some $\lambda \in [0,1]$, as copy-number proportions are non-negative. To derive $\lambda$, we substitute into Eq. (S2) which must be satisfied for consistency.

$$v = \frac{1}{F}\sum_{(x,y,m)\in\Gamma} m\cdot g_{(x,y,m)}$$

$$v = \frac{1}{F}\left[0\cdot(1-\lambda)\mu_{(x^*,y^*)} + \cdot m^*\cdot\lambda\mu_{(x^*,y^*)} + \sum_{\substack{(x,y,m)\in\Gamma \\ (x,y)\neq(x^*,y^*)}} m\cdot\mu_{(x,y)}\right]$$

Solving for $\lambda$, we obtain

$$\lambda = \frac{1}{m^*\mu_{(x^*,y^*)}}\left[vF - \sum_{\substack{(x,y,m)\in\Gamma^* \\ (x,y)\neq(x^*,y^*)}} m\cdot\mu_{(x,y)}\right].$$

Thus, any consistent genotype proportions will be fully determined by the above equations. □

Note that $\lambda$ has a natural interpretation as the the proportion of cells with copy-number state $(x,y)$ that have the mutation, thus quantifying the "split" of the single split copy-number state.

Lemma 1 gives the closed form of consistent genotypes but notes that there may not exist consistent genotype proportions for a given VAF $v$, copy-number proportions $\mu$, and SSCN genotype set $\Gamma^*$. Below, we describe two conditions that together are necessary and sufficient for the existence of consistent genotype proportions.

**Lemma 2**

Given VAF $v$, copy-number proportions $\boldsymbol{\mu}$, and an SSCN genotype set $\Gamma^*$, genotype proportions $\mathbf{g}$ such that $(\Gamma^*, \mathbf{g})$ are consistent with VAF $v$ and copy-number proportions $\boldsymbol{\mu}$ exist if and only if

1. For all copy-number states $(x, y)$ such that $\mu_{(x,y)} > 0$, there exists state $(x, y, m) \in \Gamma$ for some $m$.
2. $\lambda$ is in the range $[0, 1]$ (Eq. (S6)).

**Proof**

To prove this, we must prove both directions of this statement. First, we will prove that if condition (1) and (2) are met, then the genotype proportions given by Eq. (S5) are consistent, i.e., they satisfy Eqs. (S2) and (S4). In the proof for Lemma 1 we show that Eq. (S2) is satisfied with $\lambda \in [0, 1]$. We will show here they satisfy Eq. (S4). Consider copy-number state $(x, y)$ such that $(x, y) \neq (x^*, y^*)$. If condition (1) is met then there exists state $(x, y, m) \in \Gamma$, with $g_{(x,y,m)} = \mu_{(x,y)}$, which satisfies Eq. (S4). Next consider copy-number state $(x, y)$ such that $(x, y) = (x^*, y^*)$. Condition (1) is necessarily met by the definition of SSCN genotype set. Thus $\mu_{(x^*,y^*)} = g_{(x^*,y^*,0)} + g_{(x^*,y^*,m^*)} = (1 - \lambda)\mu_{(x^*,y^*)} + \lambda\mu_{(x^*,y^*)}$, which satisfies Eq. (S4).

Now we will prove the reverse direction, i.e., if the genotype proportions given by Eq. (S5) are consistent, then conditions (1) and (2) are met. Consider condition (1) and assume, for the sake of contradiction, the converse: for some copy-number state $(x, y)$ with $\mu_{(x,y)} > 0$, there does not exist any state $(x, y, m) \in \Gamma$. Thus $\sum_{(x,y,m)\in\Gamma} g_{(x,y,m)} = 0$ and thus the genotype proportions are not consistent. Thus we have a contradiction. Consider condition (2), and assume, for the sake of contradiction, the converse: $\lambda \notin [0, 1]$. If $\lambda \notin [0, 1]$, then either $g_{(x^*,y^*,0)} < 0$ or $g_{(x^*,y^*,m^*)} < 0$, which is not allowed by definition of genotype proportions. Thus, there do not exist consistent genotype proportions, and we have a contradiction. □

The constraint on $\lambda \in [0, 1]$ also defines the range of VAFs that are possible for a given $\boldsymbol{\mu}$ and $\Gamma^*$.

**Lemma 3**

A VAF $v$ is *feasible* for copy-number proportions $\boldsymbol{\mu}$ and an SSCN genotype set $\Gamma^*$ consistent with $\boldsymbol{\mu}$

provided $v$ is in the range $[v^-, v^+]$ where

$$v^- = \frac{1}{F} \sum_{\substack{(x,y,m)\in\Gamma^* \\ (x,y)\neq(x^*,y^*)}} m \cdot \mu_{(x,y)} \qquad \text{and} \qquad v^+ = v^- + = \frac{m^* \cdot \mu_{(x^*,y^*)}}{F}. \tag{S7}$$

**Proof**

We begin by finding $v^-$ such that

$$v^- = \min_{\mathbf{g}} \frac{1}{F} \sum_{(x,y,m)\in\Gamma^*} m \cdot g_{(x,y,m)}$$

where $\mathbf{g}$ must be consistent with $\boldsymbol{\mu}$. We separate out the genotype proportions for the split state $(x^*, y^*)$.

$$\begin{aligned} v^- &= \min_{\mathbf{g}} \frac{1}{F} \left( m^* \cdot g_{(x^*,y^*,m^*)} + 0 \cdot g_{(x^*,y^*,0)} + \sum_{\substack{(x,y,m)\in\Gamma^* \\ (x,y)\neq(x^*,y^*)}} m \cdot g_{(x,y,m)} \right) \\ &= \min_{\mathbf{g}} \frac{1}{F} \left( m^* \cdot g_{(x^*,y^*,m^*)} + \sum_{\substack{(x,y,m)\in\Gamma^* \\ (x,y)\neq(x^*,y^*,m^*)}} m \cdot g_{(x,y,m)} \right) \end{aligned}$$

Recall from Eq. (S5) that for all $(x, y) \neq (x^*, y^*)$, $g_{(x,y,m)} = \mu_{x,y}$. Thus, we have

$$\begin{aligned} v^- &= \min_{\mathbf{g}} \frac{1}{F} \left( m^* \cdot g_{(x^*,y^*,m^*)} + \sum_{\substack{(x,y,m)\in\Gamma^* \\ (x,y)\neq(x^*,y^*)}} m \cdot \mu_{(x,y)} \right) \\ &= \min_{g_{(x^*,y^*,m^*)}} \frac{1}{F} \left( m^* \cdot g_{(x^*,y^*,m^*)} + \sum_{\substack{(x,y,m)\in\Gamma^* \\ (x,y)\neq(x^*,y^*)}} m \cdot \mu_{(x,y)} \right). \end{aligned}$$

Thus $v$ is minimized when $g_{(x^*,y^*,m^*)}$ is minimized. By Eq. (S5), we have that $g_{(x^*,y^*,m^*)} = \lambda\mu_{(x,y)}$ and $\lambda \in [0, 1]$. Thus $g_{(x^*,y^*,m^*)}$ is minimized when $\lambda = 0$ and $g_{(x^*,y^*,m^*)} = 0$. Therefore,

$$v^- = \frac{1}{F} \left( \sum_{\substack{(x,y,m)\in\Gamma^* \\ (x,y)\neq(x^*,y^*)}} m \cdot \mu_{(x,y)} \right)$$

Following the same logic, we have that $v$ is maximized when $g_{(x^*,y^*,m^*)}$ is maximized, i.e., when $\lambda = 1$.

Thus, we have that

$$v^+ = \frac{1}{F}\left( m^* \cdot \mu_{(x^*,y^*)} + \sum_{\substack{(x,y,m)\in\Gamma^* \\ (x,y)\neq(x^*,y^*)}} m \cdot \mu_{(x,y)} \right) = v^- + \frac{m^* \cdot \mu_{(x^*,y^*)}}{F}$$

□

As the CCF is determined by genotype proportions, we can use Lemma 1 to characterize CCF $c$ in terms of $\lambda$.

**Lemma 4**

Given (i) copy-number proportions $\boldsymbol{\mu}$, (ii) an SSCN genotype set $\Gamma$ and (iii) the tumor purity $\rho$, the CCFs $c$ resulting from genotype proportions $\mathbf{g}$ such that $(\Gamma, \mathbf{g})$ is consistent with $\boldsymbol{\mu}$ are uniquely determined as

$$c = \frac{1}{\rho}\left[ \lambda \cdot \mu_{(x^*,y^*)} + \sum_{\substack{(x,y,m)\in\Gamma_{\mathrm{CCF}} \\ (x,y)\neq(x^*,y^*)}} \mu_{(x,y)} \right] \tag{S8}$$

for any value of the splitting parameter $\lambda \in [0, 1]$.

**Proof**

Follows from Lemma 1 and Definition 1. □

We can thus derive Main Text Theorem 1.

**Theorem 1**

Given tumor purity $\rho$, VAF $v$, copy-number proportions $\boldsymbol{\mu}$, and an SSCN genotype set $\Gamma^*$ consistent with $v$ and $\boldsymbol{\mu}$, the CCF $c$ is uniquely determined by

$$c = \frac{1}{\rho m^*}[vF - \sum_{\substack{(x,y,m)\in\Gamma_{\mathrm{CCF}} \\ (x,y)\neq(x^*,y^*)}} (m - m^*) \cdot \mu_{(x,y)}], \tag{S9}$$

where $\Gamma_{\mathrm{CCF}} = \{(x, y, m) \in \Gamma^* \mid m \geq 1\} \subseteq \Gamma^*$ is the set of genotypes containing the mutation.

**Proof**

Substituting Eq. (S5) into Eq. (S2) to obtain the following.

$$v = \frac{1}{F}\left[m^* \cdot \lambda \cdot \mu_{(x^*,y^*)} + \sum_{\substack{(x,y,m)\in\Gamma \\ (x,y)\neq(x^*,y^*)}} m \cdot \mu_{(x,y)}\right]$$

We solve this for the term $\lambda \cdot \mu_{(x^*,y^*)}$.

$$m^* \cdot \lambda \cdot \mu_{(x^*,y^*)} = Fv - \sum_{\substack{(x,y,m)\in\Gamma \\ (x,y)\neq(x^*,y^*)}} m \cdot \mu_{(x,y)}$$

$$\lambda \cdot \mu_{(x^*,y^*)} = \frac{Fv}{m^*} - \frac{1}{m^*}\sum_{\substack{(x,y,m)\in\Gamma \\ (x,y)\neq(x^*,y^*)}} m \cdot \mu_{(x,y)}$$

Substituting in for $\lambda \cdot \mu_{(x^*,y^*)}$ in Eq. (S8) yields.

$$c = \frac{1}{\rho}\left[\frac{Fv}{m^*} - \frac{1}{m^*}\sum_{\substack{(x,y,m)\in\Gamma \\ (x,y)\neq(x^*,y^*)}} m \cdot \mu_{(x,y)} + \sum_{\substack{(x,y,m)\in\Gamma_{\text{CCF}} \\ (x,y)\neq(x^*,y^*)}} \mu_{(x,y)}\right]$$

Note that $m = 0$ when $(x, y, m) \notin \Gamma_{\text{CCF}}$ and thus

$$\sum_{\substack{(x,y,m)\in\Gamma \\ (x,y)\neq(x^*,y^*)}} m \cdot \mu_{(x,y)} = \sum_{\substack{(x,y,m)\in\Gamma_{\text{CCF}} \\ (x,y)\neq(x^*,y^*)}} m \cdot \mu_{(x,y)}.$$

From here, we obtain Theorem 1.

$$c = \frac{1}{\rho}\left[\frac{Fv}{m^*} - \frac{1}{m^*}\sum_{\substack{(x,y,m)\in\Gamma_{CCF} \\ (x,y)\neq(x^*,y^*)}} m \cdot \mu_{(x,y)} + \sum_{\substack{(x,y,m)\in\Gamma_{\text{CCF}} \\ (x,y)\neq(x^*,y^*)}} \mu_{(x,y)}\right]$$

$$c = \frac{1}{\rho}\left[\frac{Fv}{m^*} - \sum_{\substack{(x,y,m)\in\Gamma_{\text{CCF}} \\ (x,y)\neq(x^*,y^*)}} \frac{m}{m^*} \cdot \mu_{(x,y)} + \sum_{\substack{(x,y,m)\in\Gamma_{\text{CCF}} \\ (x,y)\neq(x^*,y^*)}} \frac{m^*}{m^*}\mu_{(x,y)}\right]$$

$$c = \frac{1}{\rho}\left[\frac{Fv}{m^*} - \sum_{\substack{(x,y,m)\in\Gamma_{\text{CCF}}\\(x,y)\neq(x^*,y^*)}} \frac{m-m^*}{m^*}\cdot\mu_{(x,y)}\right]$$

$$c = \frac{1}{\rho m^*}\left[Fv - \sum_{\substack{(x,y,m)\in\Gamma_{\text{CCF}}\\(x,y)\neq(x^*,y^*)}} (m-m^*)\cdot\mu_{(x,y)}\right]$$

□

#### B.2 Derivation of the CCF formula under the CMM Assumption

We restate the CMM assumption given in the main text.

**Assumption**

**Constant Mutation Multiplicity (CMM).** At every SNV locus, there exists an integer $M \geq 1$ such that all genotypes at the locus have the form $(x, y, m)$ where either $m = 0$ or $m = M$.

Under this assumption, many methods approximate the CCF as

$$c \approx \frac{1}{\rho}\left(\frac{\rho N_{\text{tot}} + (1-\rho)\,2}{M}\right)\hat{v}, \quad \text{(S10)}$$

where $N_{\text{tot}}$ is the average copy number of all cancer cells. By contrast, the fractional copy number $F$ is the average copy number of all cells, tumor and normal cells alike. We have the following relationship for SNVs in autosomal chromosomes.

$$F = \rho N_{\text{tot}} + 2(1-\rho). \quad \text{(S11)}$$

We will now show that the this equation approximates the CCF, as given by Definition 1.

**Proposition 5**

Under the CMM assumption with SNV multiplicity $M$ and average copy number $N_{\text{tot}}$, variant allele frequency $\hat{v}$ and tumor purity $\rho$, the CCF is approximated as

$$c \approx \frac{1}{\rho}\left(\frac{\rho N_{\text{tot}} + (1-\rho)\,2}{M}\right)\hat{v}, \quad \text{(S12)}$$

**Proof**

We begin using Definition 1 that defines the CCF in terms of $(\Gamma, \mathbf{s}, \rho)$.

$$c = \frac{1}{\rho}\sum_{(x,y,m)\in\Gamma} g_{(x,y,m)} = \frac{1}{\rho}\sum_{\substack{(x,y,m)\in\Gamma\\ m\geq 1}} g_{(x,y,m)}.$$

Next, we use the CMM assumption, stating that for all $(x, y, m) \in \Gamma$ where $m > 0$ it holds that $m = M$.

$$c = \frac{1}{\rho} \sum_{\substack{(x,y,m)\in\Gamma \\ m\geq 1}} g_{(x,y,M)} \tag{S13}$$

We use Definition 2 that defines the VAF in terms of $(\Gamma, \mathbf{s})$.

$$v = \frac{1}{F} \sum_{\substack{(x,y,m)\in\Gamma \\ m\geq 1}} m \cdot g_{(x,y,m)}.$$

Next, we use the CMM assumption, stating that for all $(x, y, m) \in \Gamma$ where $m > 0$ it holds that $m = M$.

$$v = \frac{M}{F} \sum_{\substack{(x,y,m)\in\Gamma \\ m\geq 1}} g_{(x,y,M)}$$

Then, we rearrange terms.

$$\sum_{\substack{(x,y,m)\in\Gamma \\ m\geq 1}} g_{(x,y,M)} = \frac{vF}{M} \tag{S14}$$

We substitute (S14) in (S13), leading to

$$c = \frac{Fv}{\rho M}.$$

Finally, we use (S11), leading to

$$c = \frac{1}{\rho} \left( \frac{\rho N_{\text{tot}} + (1-\rho)\, 2}{M} \right) \hat{v}.$$

□

#### B.3 Probabilistic model for CCF

In Theorem 1, we showed that Eq. (5) is an affine transformation of CCF to VAF for SSCN genotypes. We can likewise transform a probability distribution over VAF $v$ to a probability distribution over CCF $c$ using the change-of-variables technique. In particular, if we have random variables $c$ and $v$ such that $c = \tau(v)$ where $\tau$ is a monotonic, then it holds that

$$P_c(c) = P_v(\tau^{-1}(c)) \left| \frac{d}{dy} \tau^{-1}(c) \right| . \tag{S15}$$

where $P_F$ and $P_C$ are the probability density functions with respect to $f$ and $c$ respectively. From Eq. (S10), we compute the inverse transformation $\tau^{-1}(c) = V(c)$ as

$$V(c) = \frac{c\rho m^*}{F} + \frac{1}{F} \sum_{\substack{(x,y,m)\in\Gamma_{\mathrm{CCF}} \\ (x,y)\neq(x^*,y^*)}} (m - m^*) \cdot \mu_{(x,y)}. \tag{S16}$$

Thus, given any probability distribution over $v$, we have the following probability distribution over $c$.

$$\Pr(c \mid a, t, \rho, \boldsymbol{\mu}, \Gamma) = \frac{\rho m^*}{F} \Pr(V(c) \mid a, t, \boldsymbol{\mu}, \Gamma) \tag{S17}$$

To derive $\Pr(V(c) \mid a, t, \boldsymbol{\mu}, \Gamma)$, we first apply Bayes' Theorem, giving

$$\Pr(V(c) \mid a, t, \boldsymbol{\mu}, \Gamma) \propto \Pr(V(c) \mid t, \boldsymbol{\mu}, \Gamma) \Pr(a \mid V(c), t, \boldsymbol{\mu}, \Gamma). \tag{S18}$$

We then assume that the VAF $V(c)$ is conditionally independent of the total read count $t$ given $\boldsymbol{\mu}$ and $\Gamma$ and that variant read count $v$ is conditionally independent of $\boldsymbol{\mu}$ and $\Gamma$ given the VAF $V(c)$ and total read count $t$. This yields the following posterior probability for $V(c)$.

$$\Pr(V(c) \mid a, t, \boldsymbol{\mu}, \Gamma) \propto \Pr(V(c) \mid \boldsymbol{\mu}, \Gamma) \Pr(a \mid V(c), t). \tag{S19}$$

The first component $\Pr(a \mid v, t, \boldsymbol{\mu}, \Gamma) = \Pr(a \mid v, t)$ is the likelihood of observing $a$ variant reads out of $t$ total reads, given a VAF $v$. The second component $\Pr(v \mid t, \boldsymbol{\mu}, \Gamma) = \Pr(v \mid \boldsymbol{\mu}, \Gamma)$ is the prior probability of an SNV having VAF $v$ given the copy-number proportions $\boldsymbol{\mu}$ and genotype set $\Gamma$. In Lemma 3 in Appendix B.1, we show that not all VAFs are feasible for a given $\boldsymbol{\mu}$ and $\Gamma$. For example, $0.5$ is an upper bound for VAFs $v$ for heterozygous mutations in diploid regions. We also define the feasible range $[v^-, v^+]$ for VAFs. Thus, $P(v \mid \boldsymbol{\mu}, \Gamma)$ has support only on $[v^-, v^+]$. Any probability distribution that has this property may be used as a prior for $v$. DeCifer uses either the binomial or beta-binomial model for the likelihood of the variant read count $a$ given total read count $d$ and VAF $v(c)$ as is standard in the field. For the prior, DeCiFer uses a uniform prior over the feasible range of VAFs, $P(v(c) \mid \boldsymbol{\mu}, \Gamma) \propto \mathbf{1}_{v(c)\in[v^-,v^+]}$. Thus, DeCiFer uses the following posterior distribution over the CCF $c$.

$$P(c \mid a, t, \rho, \boldsymbol{\mu}, \Gamma) = \frac{1}{Z} \mathbf{1}_{V(c)\in[v^-,v^+]}\ B(a \mid V(c), t, \ldots) \tag{S20}$$

where $B$ is the binomial or beta-binomial distribution and Z is a normalization constant. In Appendix B.7 we describe how we estimate parameters for the beta-binomial distribution from sequence data.

**DCF** Using a similar procedure, we may obtain a posterior probability distribution over DCFs $d$. Here, the inverse transformation $V(c)$ is replaced with

$$V(d) = \frac{d\rho m^*}{F} + \frac{1}{F} \sum_{\substack{(x,y,m)\in\Gamma_{\mathrm{DCF}} \\ (x,y)\neq(x^*,y^*)}} (m - m^*)\mu_{(x,y)} \tag{S21}$$

Note that $\Gamma_{\mathrm{DCF}}$ is the set of genotypes in genotype tree $T_{\Gamma^*}$ that are descendants of the state $(x^*, y^*, m^*)$. Thus $V(d)$ depends on both the genotype set $\Gamma$ and the genotype tree $T_\Gamma$. We thus have

$$\Pr(d \mid a, t, \rho, \boldsymbol{\mu}, \Gamma, T_\Gamma) = \frac{\rho m^*}{F} \Pr(V(d) \mid a, t, \boldsymbol{\mu}, \Gamma). \tag{S22}$$

The probability model for VAFs is unchanged. Thus, using a uniform prior and binomial or beta-binomial likelihood, we have

$$P(d \mid a, t, \rho, \boldsymbol{\mu}, \Gamma, T_\Gamma) = \frac{1}{Z} \mathbf{1}_{V(d) \in [v^-, v^+]}\, B(a \mid V(d), t, \ldots). \tag{S23}$$

#### B.4 Mutation clustering and genotype selection

We formalize the theoretical problem of SNV clustering and selection when VAFs $v$ are observed rather than read counts $a$, $t$. In this case, let $\mathcal{G}_i$ be the set composed of pairs $(\Gamma_i^*, T_i)$ of SSCN genotype sets and genotype trees that are consistent with $\boldsymbol{\mu}$. Note that when $v$ and $\mu$ are observed, the DCF $d$ is fully determined by a pair $(\Gamma_i^*, T_i)$ and the observed copy-number proportions $\boldsymbol{\mu}$ and purity $\rho$ (see 2 Eq. (10) in Theorem 2). Let $s_i \in \{1, \ldots |\mathcal{G}_i|\}$ be a selection of a genotype set and genotype tree pair for SNV $i$ This leads to the following problem.

**Problem 2** (Mutation Clustering and Genotype Selection (MCGS))**.** *Given (i) an integer $k > 0$, and for each SNV i, (ii) a set $\mathcal{G}_i$ of pairs of genotype sets and trees; find (i) a set $\mathcal{D}$ of size $k$ of DCF values and, for each SNV i, select (ii) $s_i \in \{1, \ldots |\mathcal{G}_i|\}$ such that $d_i \in \mathcal{D}$ where $d_i$ is the DCF determined by $s_i$.*

The MCGS problem is an instance of the well-known Hitting Set problem, which is known to be equivalent to the Set Cover problem and is NP-complete.

#### B.5 DeCiFer algorithm

DeCiFer aims to solve the PMCGS problem, which optimizes the following function.

$$D^*, \mathbf{s}^*, \mathbf{z}^* = \operatorname*{argmax}_{D, \mathbf{s}, \mathbf{z}} \prod_{i=1}^{n} \prod_{\ell=1}^{p} \Pr(d_{z_i, \ell} \mid a_{i,\ell}, t_{i,\ell}, \boldsymbol{\mu}_{i,\ell}, \Gamma_{i,s_i}, T_{i,s_i}), \tag{S24}$$

We use a coordinate ascent approach, where we alternatively optimize (i) cluster DCF values $D$ and (ii) SNV cluster $\mathbf{s}$ and genotype tree assignments $\mathbf{z}$. We begin by randomly initializing $D^{(0)}$ following $k$ draws from a symmetric Dirichlet distribution. Note that given $D$, genotype tree and cluster assignments $s_i$ and $z_i$ for an individual SNV are conditionally independent of the cluster and genotype set assignments of other SNVs. Thus, we optimize separately for each SNV $i$,

$$s_i^{(q)}, z_i^{(q)} \mid D^{(q-1)} = \operatorname*{argmax}_{\substack{s \in \{1, \ldots, |\mathcal{G}_i|\}, \\ z \in \{1, \ldots, k\}}} \prod_{\ell=1}^{p} \Pr(d_{z,\ell}^{(q-1)} \mid a_{i,\ell}, t_{i,\ell}, \boldsymbol{\mu}_{i,\ell}, \Gamma_{i,s}, T_{i,s}), \tag{S25}$$

by considering all possible cluster and genotype set assignments. We then find the optimal cluster cell fraction values $D^{(q)} \in [0,1]^{k \times p}$. Given $\mathbf{s}^{(q)}$ and $\mathbf{z}^{(q)}$, a DCF value $d_{\ell,j}$ in sample $\ell$ for cluster $j$ is conditionally independent of all other DCF values. Thus, we find

$$d_{\ell,j}^{(q)} \mid \mathbf{s}^{(q)}, \mathbf{z}^{(q)} = \operatorname*{argmax}_{d \in [0,1]} \prod_{i; z_i^{(q)} = j} \Pr(d \mid a_{i,\ell}, t_{i,\ell}, \boldsymbol{\mu}_{i,\ell}, \Gamma_{i,s_i^{(q)}}, T_{i,s_i^{(q)}}). \tag{S26}$$

DeCiFer optimizes Eq. (S26) using Brent's algorithm to find a minimum in the range $[0, 1]$, as implemented in the `minimize_scalar` method of the SciPy optimize package[62]. DeCiFer terminates upon convergence when cluster and genotype set assignments do not change between iterations, which indicates that the objective function does not change. Coordinate ascent is not guaranteed to converge to the global maximum. Thus, we use multiple restarts with different initializations.

#### B.6 Model selection

The number $k$ of clusters is unknown, and thus we estimate it using a model-selection criterion. Specifically, we consider two different classes of clusters. The first class includes $p + 2$ fixed clusters, which we assume to be potentially present in every tumor. The second class includes a variable number of clusters, which is estimated using a model-selection criterion and whose cell fractions are estimated by DeCiFer. We describe the details of these clusters in the remaining of the section.

For the first class of clusters, we introduce $p + 2$ fixed clusters, composed of three different groups. First, we include a 'truncal' cluster, such that the cell fraction values $\mathbf{d}_T = [\rho_\ell]_{l\in[1,\ldots,p]}$ for this cluster are fixed to the purity of the samples. This is motivated by the common observation that in many tumors, there are a large number of SNVs that are present in all cells. Second, we include an 'absent' cluster, such that the cell fraction values $\mathbf{d}_0 = [0]_{l\in[1,\ldots,p]}$ for this cluster are fixed to zero. This cluster is necessary to guarantee feasiblity of solutions during optimization. That is, during early optimization steps, there exist mutations such that none of the cell fraction values are feasible. As such, the posterior probability for all cell fraction values would be zero, and the coordinate-ascent algorithm would be stuck. However, for SSCN genotype trees meeting evolutionary constraints, there always exists a genotype tree for which $d = 0$ is a feasible cell fraction value. Thus, using an 'absent' cluster guarantees non-zero objective functions during optimization. Moreover, this 'absent' cluster may capture false positive variant calls in real data. The cell fraction values for the 'truncal' and 'absent' clusters do not change during optimization. Last, we include $p$ clusters which represent SNVs specific to each sample. This is motivated by the observations that frequently every sample has a certain number of unique SNVs. For a sample specific cluster in sample $\ell$, we fix the cell fraction value of all other samples to be 0. We initialize $d_\ell = \rho_\ell$ to be the sample purity, but this value may change during optimization.

For the second class of clusters, we choose their number using a model-selection criterion based on the standard elbow method[63]. Specifically, we consider an increasing number $k$ of clusters, starting from the minimum of $p + 2$ as described above until a given maximum number of clusters, and we apply DeCiFer to each of these values. As such, we evaluate how the objective value computed by DeCiFer changes with varying the number of clusters and we choose $k$ as the elbow of the function, i.e., the value which improves the objective with respect to lower values but is not substantially improved by subsequent increases. Formally, we evaluate such improvements by using previous elbow methods[32,64], based on comparing the differences between the left and right derivatives of the objective function in each point.

##### B.7 Estimating beta-binomial parameters

Read counts from bulk DNA sequencing are often affected by overdispersion, as noted in previous studies[38]. We account for such overdispersion in DeCiFer by using a beta-binomial generative model similarly to the one proposed for PyClone[38]. Specifically, we model the variant read count $a$ with a Beta-Binomial distribution parameterized by mean $v$ and precision $\mathrm{s}$. However, in contrast to previous approaches[38] that define a prior distribution of $\mathrm{s}$, in this work we estimate the value of $\mathrm{s}$ using germline single-nucleotide polymorphisms (SNPs). In particular, under the assumption that read counts from somatic SNVs or germline SNPs are affected by the same level of overdispersion (i.e., are drawn from Beta-Binomial distributions with the same parameters) we choose $\mathrm{s}$ to best fit germline SNPs with the same Beta-Binomial distributions parameterized by the mean $v$ and $\mathrm{s}$. Note that we know the value of $v$ for germline SNPs, as this value is provided by the inferred allele-specific copy numbers: specifically, $v$ corresponds to the proportion of allele-specific copy number, namely the B-allele frequency (BAF)[32]. Moreover, we only consider SNPs with BAF $= 0.5$ or $\not\approx 0.5$ to SNPs for which the computation of BAF is challenging[32]. We thus use the inferred value $s$ and we substitute the likelihood $\Pr(a \mid v, d, \mu, \Gamma) = \text{Binomial}(a \mid v, d)$ with $\Pr(a \mid v, d, \mu, \Gamma, s) = \text{Beta-Binomial}(a \mid v, d, s)$. Note that DeCiFer can use either generative model, according to different kinds of sequencing data.

##### B.8 Simulated data analysis details

**DeCiFer** On simulated data, DeCiFer was run using the following parameters:

| Parameter | Value |
|---|---|
| Likelihood model | Binomial |
| Min # of clusters | 2 |
| Max # of clusters | 15 |
| # of restarts | 10 |
| Max # of iterations | 50 |
| Elbow parameter | .2 |
| # of threads | 30 |

**PyClone** PyClone does not consider subclonal copy-number aberrations and instead takes as input integer allele-specific copy number $N_{maj}, N_{min}$ for each SNV. As the simulated data contains subclonal copy-number aberrations, we used two approaches to provide input for PyClone. In the first, we estimated the average allele-speciic copy number for each SNV and rounded to the nearest integer value. That is,

$$N_{maj} = \lceil \max\left(\sum_{x,y} x \cdot \mu_{x,y}, \sum_{x,y} y \cdot \mu_{x,y}\right) \rfloor \tag{S27}$$

and

$$N_{min} = \lceil \min\left(\sum_{x,y} x \cdot \mu_{x,y}, \sum_{x,y} y \cdot \mu_{x,y}\right) \rfloor. \tag{S28}$$

In the second approach, we used a method to compute CCFs that accounts for subclonal CNAs[41], and used PyClone to cluster these CCFs based on scaled input VAFs. That is, we compute $\hat{c}$ using the method outlined by Dentro et al.[41]. Then as input to PyClone, we use $N_{maj} = N_{min} = 1$, the number of total reads $t$ is the simulated total number of reads and the number of variant reads is $a = \lceil \hat{c} \cdot t \rfloor$.

**PhyloWGS** PhyloWGS was run using default parameters.

##### B.9 Bioinformatic analysis of metastatic prostate cancer patients

DeCiFer requires two inputs for every sample in each patient: (1) sequencing read counts for SNVs and (2) allele-specific copy numbers for genomic regions harboring SNVs. We considered SNVs previously identified in these samples by Zaccaria & Raphael[32] using Varscan 2 (v2.3.9)[65]. To minimize false positives in SNV calls, we selected high confidence SNVs using the p-value computed by Varscan 2 ($p < 10^{-4}$ in at least one sample). Varscan 2 analyzes each sample independently and does not report SNVs without a sufficiently high number of variant reads. As such, we used BCFtools (v1.9) to obtain read counts across all samples for the set of SNVs called by Varscan 2 in at least one sample. We used the allele-specific copy numbers previously inferred by HATCHet for the same samples[32]. To enable the efficient enumeration of all possible state trees and to limit potential errors in regions with high copy numbers, we excluded any genomic region with allele-specific copy numbers $(x, y)$ where $\max(x, y) > 4$ and $\min(x, y) > 2$. We applied DeCiFer to all patients using the default values of all parameters and by considering a beta-binomial generative model with precision parameter $s \approx 200$, as estimated from germline SNPs.

#### C Supplementary Results

##### C.1 Analysis of DCF in prostate cancer patients

In this study, we specifically focus on three categories of SNVs that have been classified differently using DCFs inferred by DeCifer as compared to CMM CCFs: (1) SNVs classified as truncal by DeCiFer and as subclonal by the CMM CCFs, (2) SNVs classified as subtruncal by DeCiFer and clonal by the CMM CCFs, and (3) SNVs classified as truncal or subtruncal by DeCiFer and absent by the CMM CCFs. Interestingly, we observe that the patients with the highest fractions of SNVs with different classifications (A29, A31, and A32) are those for which whole-genome duplications (WGDs) have been previously predicted[32]. This is expected as WGDs double the entire genome content, increasing the copy numbers of every genomic region and yielding more combinations of different mutation multiplicities that may violate the CMM assumption (Figure S5b–c).

##### C.2 Identification of subtruncal SNVs in prostate cancer patients

DeCiFer classifies ~12,000 SNVs as subtruncal across all samples that are classified as clonal using CMM CCFs (Figure S5a). In individual patients, such SNVs correspond to between 3–41% of the total number of SNVs in different patients (Figure S5c). As in Figure 4, we see that such classification differences may result in unlikely phylogenetic reconstructions with extensive homoplasy. For example, 167 SNVs in chromosome 7 of prostate cancer patient A24 form four distinct groups with different VAFs in sample A24-A. In this genomic region, there are two groups of cancer cells with different copy numbers (Figure S6a,b): 27% of cancer cells have an LOH with amplifications (i.e., allele-specific copy numbers $\{4, 0\}$), while the remaining cancer cells are diploid (i.e., allele-specific copy numbers $\{1, 1\}$). Using the CMM CCFs, these SNVs form two clusters (Figure S6c): one subclonal cluster composed of 30 SNVs, and one clonal cluster composed of the remaining 137 SNVs. Based on the CMM assumption, all the clonal SNVs are inferred to have a constant SNV multiplicity of either 1 or 2. However, this solution is biologically unlikely: If an SNV was present in the cell where the amplification occurred, then the cell would have 4 copies of the SNV. If there are only 1 or 2 copies, this implies that for 137 SNVs, there is either a homoplasy event in the diploid cell or back mutation(s) in the tetraploid cells. DeCiFer identifies that one group of 60 SNVs (light and dark blue) belonging to a subtruncal cluster (CCF$\approx 0.7$) while the remaining SNVs (light and dark grey) belong to a truncal cluster (Figure S6d). Moreover, DeCiFer explains the different VAFs for groups of SNVs in the same cluster due to the presence of different mutation multiplicities in different clones for mutations on the retained allele and the loss of the mutations on the other lost allele. The solution of DeCiFer thus results in a realistic reconstruction of tumor evolution, where the differences in mutation multiplicities that contradict the CMM assumption are consistent with the amplification of the related allele.
